# Single-Cell Atlas of Renal Cell Carcinoma Brain Metastasis Uncovers Mechanisms of Immune Dysfunction and Resistance

**DOI:** 10.64898/2026.05.06.722652

**Authors:** Mostafa I.H. Ali, Zeynep Feyza Akpinar, Jose A Ovando-Ricardez, Anna K. Casasent, Truong Nguyen Anh Lam, Jerome Lin, Narmina Khanmammadova, Patrick K. Reville, David J. H. Shih, Adeboye O. Osunkoya, Lisa M. Norberg, Tuan M. Tran, Jianzhuo Li, Anh G. Hoang, Sahin Hanalioglu, Mehmet Asim Bilen, Frederick Lang, Jason T. Huse, Nicholas Navin, Merve Hasanov, Eric Jonasch, Elshad Hasanov

**Affiliations:** Division of Medical Oncology, Department of Internal Medicine, The Ohio State University Comprehensive Cancer Center, Columbus, OH, USA; Pelotonia Institute for Immuno-Oncology, The Ohio State University Comprehensive Cancer Center, Columbus, OH, USA; Department of Hematopoietic Biology & Malignancy, The University of Texas MD Anderson Cancer Center, Houston, TX, USA; Department of Genitourinary Medical Oncology, Division of Cancer Medicine, The University of Texas MD Anderson Cancer Center, Houston, TX, USA; Department of Systems Biology, Division of Discovery Science, The University of Texas MD Anderson Cancer Center, Houston, TX, USA; Department of Medicine, Division of Hematology/Oncology, Nuvance Health, Norwalk, CT; School of Biomedical Sciences, Li Ka Shing Faculty of Medicine, The University of Hong Kong, Hong Kong SAR, China; Departments of Pathology and Urology, Emory University School of Medicine, Atlanta, GA, USA; Department of Pathology, The University of Texas MD Anderson Cancer Center, Houston, TX, USA; Department of Neurosurgery, Hacettepe University Faculty of Medicine, Ankara, Turkey; Department of Hematology and Medical Oncology, Emory University School of Medicine, Atlanta, GA, USA; Department of Neurosurgery, The University of Texas MD Anderson Cancer Center, Houston, Texas

## Abstract

Brain metastasis (BM) in renal cell carcinoma (RCC) remains poorly understood and often resistant to immune checkpoint inhibitors. We generated a large single-nucleus RNA-seq data of RCC BM, profiling 14 BM samples alongside matched extracranial metastases and primary tumors. Tumor cells in BM displayed neuronal infiltration, neural-like adaptation, and marked remodeling of the microenvironment, including expansion of immunosuppressive myeloid cells and depletion of antigen-presenting dendritic cells. Tumor, immune, and stromal cells exhibited metabolic rewiring characterized by fatty-acid metabolism, oxidative phosphorylation, and MYC-driven programs. CD8⁺ T cells showed terminal exhaustion and impaired proliferative capacity, and tertiary lymphoid structures were absent. Spatial profiling of 12 BM samples (13,128 cells) validated key cellular interactions, while ligand–receptor analysis revealed immunoregulatory circuits between tumor, stromal, and immune cells. These findings define BM-specific adaptations that promote immune evasion and resistance, revealing therapeutic vulnerabilities in RCC BM.

**SIGNIFICANCE:** Single-nucleus RNA-sequencing profiling reveals tumor, immune, and metabolic adaptations in renal cell carcinoma (RCC) brain metastases, including neuroglial remodelling and immunosuppressive niche formation. These findings identify immune evasion mechanisms that could contribute to therapeutic resistance, providing new avenues for site-specific therapeutic interventions to improve treatment efficacy and outcomes in patients with RCC BM.

## INTRODUCTION

In the United States (US), recent epidemiological data reports about 80,980 new kidney and renal pelvis cancer diagnoses and 14,510 deaths per year (1). Renal cell carcinoma (RCC) accounts for approximately 85-90% of these cases (2) and brain metastases are occur in approximately 10%-12% of patients with advanced RCC (3). Primary RCC is characterized by substantial intratumoral heterogeneity, which contributes to disease progression and influences tumor-immune dynamics across metastatic sites(4–6). Systemic therapies for metastatic RCC have advanced over the past two decades with the development of targeted agents and the introduction of immune checkpoint inhibitors (ICIs) (7). However, the presence of BM is associated with a markedly poor prognosis and reduced overall survival, even in the era of immune checkpoint inhibitors (ICIs). (8), suggesting immunosuppressive mechanisms in BM (3,9–12). In a phase 2 study of nivolumab in patients with RCC BM, the median progression-free survival was 4.8 months, with a 10% objective response rate. Similarly, patients with avelumab plus axitinib treatment had 4.9 months of progression-free survival. Overall survivals has been reported to be ranging from 6 to 16 months (3,8,10). In addition to the known role of the blood-brain barrier, this also reflects unique biological features of the brain tumor microenvironment (TME), such as immune modulation, and distinct metabolic constraints that influence site-specific tumor evolution and treatment resistance (13–17). Previous single-cell and spatial profiling of the BM TME across other cancer types has uncovered transcriptional programs involving immune suppression (18,19). Yet, how RCC cells and the surrounding stromal and immune compartments transcriptionally and functionally adapt to the brain microenvironment remains poorly understood. Moreover, the mechanisms of immune evasion in RCC BM and their potential for therapeutic targeting have not been comprehensively explored. To address these gaps, we performed single-nucleus RNA sequencing (snRNA-seq) on matched primary kidney tumors (KTs), extracranial metastases (ECM), and BM from RCC patients. Additionally, we validated our key findings using CosMx Spatial Molecular Imaging (SMI) to better characterize the key ligand-receptor and spatial interactions within RCC brain TME cells. These insights reveal the multifaceted adaptations within the BM TME and provide a foundation for developing novel therapeutic strategies to overcome resistance and improve RCC patient outcomes.

## RESULTS

### Neuroglial Cell-Driven Remodelling of the RCC Brain Metastasis TME

To dissect the cellular heterogeneity of RCC across its primary and metastatic sites, we performed snRNA-seq on 27 freshly frozen tumor samples, comprising 14 BM, 8 matched KT, and 5 matched ECM (**Fig. 1A; Supplementary Fig. S1A-B**). Following tissue dissociation, library preparation, snRNA-seq data generation, and quality control filtering, we obtained transcriptomic profiles from 184,037 high-quality nuclei across all compartments. Using unsupervised graph-based clustering of all cells, we identified 9 major cell types across different tumor sites and patient samples (**Fig. 1B**), which were subsequently annotated using canonical lineage-defining markers from Cellmarker and Human Protein Atlas databases (20,21). Immune cell types included T cells (*THEMIS, CD96, CD247, TOX*), B cells (*IGHGP, IGKC, IGHG3, IGHG1*), and myeloid cells (*MSR1, MS4A6A, CD163, F13A1*). Stromal cell types comprised endothelial cells (*VWF, FLT1, CDH13*), fibroblasts (*COL1A2, LAMA2, FBN1, COL3A1, DCN*), and pericytes (*PDGFRB, RGS5*). Other defined cell types were tumor cells (*CA9, EGLN3, VEGFA, ANGPTL4, PAX8*), neuroglial (NG) cells (*RNF220, CNTN2, CTNNA2, HAPLN2, CSMD2*), and renal tubular cells (*PAX8*, *ATP6V0D2, SLC26A7, ATP6V0A4*) (**Fig. 1C; Supplementary Fig. S1C; Table S1**). Using spatial transcriptomics, we were able to validate 7 of our 9 molecular subtypes. The two cell types that were not validated NG and RTC due to rarity, organ representation, and technical features such as probe constrains (**Supplementary Fig. S1D-E; Table S2**). Subsequently, we compared the quantitative distribution of cell types across BM, ECM, and KT samples from snRNAseq data (**Fig. 1D; Table S3**). This analysis revealed that tumor cells constituted the most abundant population, consistently dominating the cellular landscape in KT, ECM, and BM TMEs. Myeloid cells represented the second most enriched cell type across all patients and tumor sites, while other immune and stromal populations, including T cells, B cells, and fibroblasts, exhibited greater variability in abundance. Each patient exhibited a distinct cellular composition, highlighting the inter-patient heterogeneity of the TME (**Supplementary Fig. S1F**). Selective enrichment of NG cells was observed in the BM TME (P <0.005), with minimal presence in KT and ECM specimens **(Fig. 1E; Supplementary Fig. S1G; Table S4**). Unsupervised analysis of NG cells identified 2 transcriptionally distinct clusters. One cluster is aligned with an oligodendrocyte/schwann cell identity, enriched for genes such as *PDE4B, PPP2R2B* and *RNF220*, which are essential for myelination and oligodendrocyte/schwann cell differentiation. The second NG cell cluster displayed a neuronal-like transcriptional profile, with elevated expression of *NLGN1, PD4ED* and *RORA*, which are associated with synaptic organization, neuronal signalling, and neurodevelopment (**Fig. 1F; Supplementary Fig. S1H; Table S5**).

**Figure 1.**
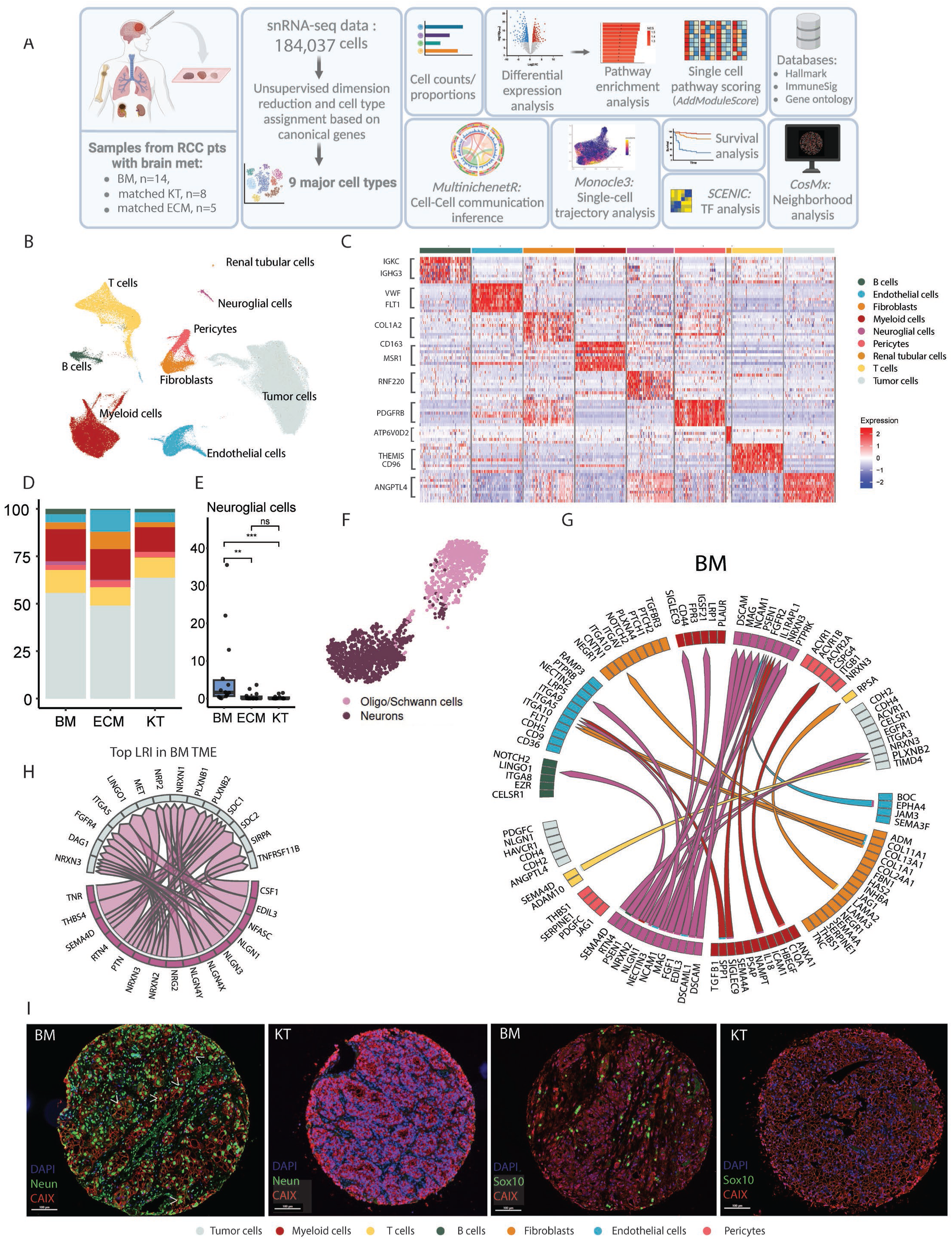
Neuroglial Cell-Driven Remodeling of the RCC Brain Metastasis TME. **(A)** Schematic overview of the study design. Single-nucleus RNA-seq (snRNA-seq) was performed on 27 freshly frozen tumor samples, including 14 brain metastases (BM), 8 matched primary kidney tumor (KT), and 5 matched extracranial metastases (ECM), to profile the renal cell carcinoma (RCC) tumor microenvironment (TME) across disease compartments (Created with BioRender.com) **(B)** UMAP representation of 184,037 nuclei after integration and unsupervised clustering, showing 9 major cell types within the TMEs of the different compartments, including tumor cells, T cells, B cells, myeloid cells, fibroblasts, endothelial cells, pericytes, renal tubular cells, and neuroglial (NG) cells. **(C)** Heatmap showing scaled, normalized expression of the top 10 differentially expressed genes per major cell type, determined using a two-sided Wilcoxon rank-sum test with Bonferroni correction for multiple testing (adjusted *P* < 0.05). **(D)** Stacked bar plot showing relative cell type proportions across all compartments (KT, ECM, and BM) highlighting tumor cells as the most abundant population, followed by myeloid cells. **(E)** Boxplot displaying the relative abundance of NG cells across the 3 anatomical compartments: BM, KT, and ECM. Statistical significance was assessed using a two-sided Wilcoxon rank-sum test. Boxplots depict the median (canter line), interquartile range (box limits), and whiskers extending up to 1.5x the interquartile range. **(F)** UMAP representation showing sub clustering of NG cells, revealing 2 transcriptionally distinct NG populations. These populations exhibit unique gene expression profiles, highlighting the heterogeneity within the NG compartment. **(G)** Circos plot illustrating intra-lineage communication among NG cells within the RCC BM TME, highlighting the top prioritized ligand-receptor interactions driving NG self-signalling. **(H)** Circos plot depicting the top prioritized ligand–receptor interactions between neuroglial cells and tumor cells within the RCC brain metastasis tumor microenvironment (TME). The connections highlight major axes of cell–cell communication that may drive neuroglial– tumor crosstalk. **(I)** Immunofluorescence staining for NeuN and SOX10 reveals strong expression of NG markers in tumor cells from RCC BM, whereas primary KT cells lack detectable expression, supporting a brain-specific neural-like phenotypic shift.

To investigate differential intercellular communication across tumor niches, a prioritization-based ligand–receptor interaction analysis was performed (22). The analysis revealed a distinct NG cell-dominated ligand-receptor interactions (LRIs) in BM (**Fig. 1G; Supplementary Fig. S1I-J**). We identified multiple interactions originating from NG cells toward other NG cells, as well as myeloid cells, B cells, endothelial cells, and tumor cells. Several receptors, including MAG (23), NCAM1 (24), NRXN3 (25,26), and FGFR2 (27) were involved in LRI among NG cells in the BM TME. These interactions are known to support neuronal differentiation, development, plasticity, and maintenance (23–27). Furthermore, an interaction involving NRXN2-SIGLEC9 between NG cells and myeloid cells was identified, indicating a potential role for NG cells in modulating the immune microenvironment (28). Among the top prioritized LRI between NG cells and tumor cells, interactions such as NRXN3-FGFR4, SEMA4D-PLXNB1, and SEMA4D-MET emerged; these are known to promote tumor proliferation, migration, invasion, and angiogenesis, thereby contributing to tumor growth and metastatic potential (29–31) (**Fig. 1H; Supplementary Fig. S1K**). We also observed NLGN-NRXN interactions between tumor and NG cells. These synaptic adhesion molecules form trans-synaptic complexes that are essential for synapse formation and neuronal communication (32). The presence of this interaction raises the possibility that RCC tumor cells may adopt a neuronal-like phenotype within the brain TME. To validate the presence of neuronal and oligodendrocyte cells in the BM TME and explore the neuronal differentiation of tumor cells, we performed multispectral imaging for the neuronal cell marker NeuN, the oligodendrocyte marker SOX10 and RCC tumor cell marker CAIX (**Fig. 1I**). Both neuronal and oligodendrocyte cells were significantly more abundant in BMs than in primary KTs. Notably, CAIX-positive tumor cells in the BM also showed strong expression of NeuN, a feature absent in primary KT samples, suggesting a brain-specific shift toward neuronal phenotype (**Fig. 1I**). To further support this observation at the transcriptomic level, we evaluated the expression of neuronal lineage and synaptic development genes across compartments. Pseudobulk analysis revealed increased expression of SOX2 (33), NEUROG2(34), NEUROG1(35), NPY, SYP, DCX, SLIT1, and SHANK3(36) in tumor cells from the BM TME compared to those from KT and ECM samples (**Supplementary Fig. S1L**). Together, these findings reveal a neuroglial remodelling within the RCC BM TME and suggest tumor cells brain-specific tumor evolution, neuronal adaptation and potential therapeutic resistance.

### Distinct Metabolic and Inflammatory State of RCC Tumor Cells in the Brain Metastasis Niche

To explore how tumor cell phenotypes are shaped by anatomical location, we performed a focused analysis of tumor cell transcriptional profiles across KT, ECM, and BM samples. Unsupervised clustering of 106,149 tumor cells, including 44,797 from BM, 15,449 from ECM, and 45,903 from KT, revealed marked inter-patient variability, with tumor cells clustering according to their patient of origin (**Fig. 2A**). Moreover, we observed distinct clustering among samples derived from the same patient but originating from different metastatic sites (**Fig. 2A-B**). Together, these findings highlight the role of the local microenvironment in shaping tumor cell identity and transcriptional state. To gain deeper insight into the transcriptional programs underlying compartment-specific differences, we performed differential gene expression analysis (DGEA) (**Fig. 2C; Table S6**). Comparisons of tumor cells from BM with those from the KT and ECM revealed a distinct brain-specific transcriptional signature enriched for metabolic reprogramming, stress adaptation, and inflammatory regulation. Specifically, when compared to KT tumor cells, BM tumor cells demonstrated upregulation of the MYC Targets V1 (*LDHA, PPIA, VDAC1*), Hypoxia signalling (*LOX, SERPINE1, GAPDH, LDHA, PGF*), mTORC1 (*DDIT4, SLC2A3, ACLY, HMGCS1, ENO1, PGM1, TPI1, GAPDH*), Oxidative Phosphorylation (OXPHOS) (*COX7C, COX5B*), Reactive Oxygen Species (ROS) pathway (GPX3, GPX4), complement system (*SERPING1, CLU*) and TNF-alpha signalling (*EGR1, SQSTM1, FOS*). In contrast, KT tumor cells exhibited enrichment of Angiogenesis (*FGFR1*), Interferon Alpha Response (SAMD9L, IFI44L, TAP1, IRF1, IL15), and Mitotic Spindle (*RABGAP1, CDK5RAP2*) **(Fig. 2C–D, Supplementary Fig. S2C; Table S6-7)**. Moreover, when compared to ECM tumor cells, BM tumor cells showed similar pathways upregulated in BM including, complement system, coagulation, OXPHOS, MYC Targets V1, ROS, mTORC1 and Hypoxia. Additionally, ECM vs. BM comparison showed elevated expression of genes associated with Epithelial-Mesenchymal Transition (EMT) (*LOX, SERPINE1*) and Fatty Acid Metabolism (*ACSL1, SLC22A5*) in BM tumor cells **(Supplementary Fig. S2A-B,D; Table S8-9)**.

**Figure 2.**
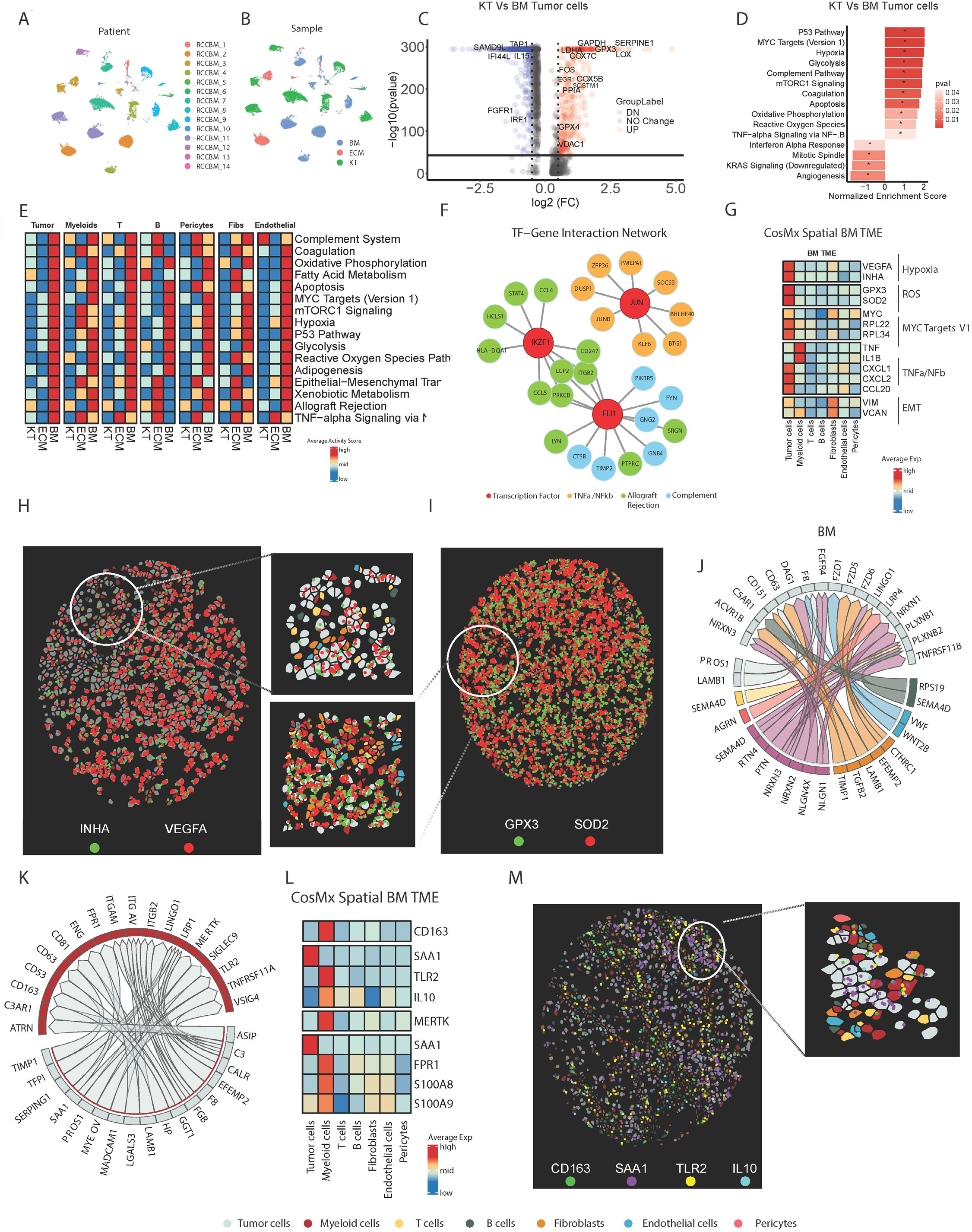
Distinct Metabolic and Inflammatory State of RCC Tumor Cells in the Brain Metastasis Niche. **(A-B)** UMAP representation of 106,149 tumor cells derived from snRNA-seq data, highlighting inter- and intra-patient heterogeneity, coloured by patient ID (left) and sample origin (right). **(C)** Volcano plot displaying differential gene expression between tumor cells from primary kidney tumors (KT) and brain metastases (BM) tumor cells. Key upregulated and downregulated gene significance determined by two-sided Wilcoxon rank-sum test. **(D)** Statistically significant Hallmark pathway enrichment analysis (FGSEA) comparing tumor cells from KT and BM. Selected pathways are shown, ordered by normalized enrichment score. **(E)** Heatmap Addmodulus activity score for Hallmark pathway of major cell types and anatomical compartments, illustrating compartment-specific biological adaptations. **(F)** Transcription factor (TF)-gene regulatory network highlighting BM-enriched TF regulons identified by SCENIC analysis. TFs (red) and their target genes (green, blue, yellow). **(G)** Pseudobulk expression heatmap of selected representative genes from top Hallmark pathways (hypoxia, MYC Targets V1, TNF-α/NF-κB signaling, EMT) across major cell types profiled by spatial transcriptomics. **(H-I)** Spatial FOVs showing colocalization of ROS representative molecules and hypoxia-related molecules within RCC BM, supporting coordinated metabolic and transcriptional adaptation. **(J)** Circos plot showing ligand–receptor interactions between immune cells, stromal cells, NG cells, and RCC tumor cells within the brain metastasis niche. tumor cells serve as the dominant receivers of TME-derived signals, underscoring major axes of tumor –TME crosstalk.**(K)** Circos plot showing ligand–receptor interactions between RCC tumor cells and myeloid cells within the brain metastasis niche. **(L)** Heatmap showing pseudobulk expression levels of key immunoregulatory molecules across major cell types within the BM TME, derived from spatial transcriptomic data. CD163 and MERTK mark immunosuppressive macrophages, while SAA1 and TLR2 highlight tumor-myeloid interactions, and IL10 represents a downstream immunosuppressive mediator. **(M)** Spatial FOVs showing colocalization of immunoregulatory axes, including CD163, SAA1, TLR2, and IL10, highlighting tumor-microglia proximity, supporting functional immune modulatory crosstalk via SAA1-TLR2-IL10 and SAA1-FPR1-S100A8/9 signaling axes.

Building on these observations, we next explored whether the enriched pathways identified in BM tumor cells were unique for tumor cells or shared across other BM TME cell types. We calculated the single-nucleus Hallmark pathway scores to compare pathway activity across the three disease compartments: BM, ECM and KT (**Supplementary Fig. S2E**). The heatmap demonstrates differential activity of Hallmark pathways across tumor and microenvironmental cell populations in KT, ECM, and BM. Compared to their counterparts in KT and ECM, BM tumor cells and non-malignant tumor microenvironment (TME) populations exhibited higher enrichment of inflammatory (complement system, TNF-α signalling, allograft rejection), metabolic (oxidative phosphorylation, fatty acid metabolism, adipogenesis, glycolysis), and stress response (p53, ROS, hypoxia, mTORC1, MYC) pathways. These findings indicate that the metabolic and signalling reprogramming observed in RCC BM is not restricted to tumor cells alone but represents a coordinated adaptation across the entire TME (**Fig. 2E**). Notably, these pathways enhance metastatic potential of tumor cells by promoting plasticity, stress resistance, immune evasion, and inflammatory microenvironment remodelling for invasion, colonization, and therapy resistance (37–43).

We further assessed Hallmark pathways activity across cell types within RCC BM TME (**Supplementary Fig. S2F**). Tumor cells exhibited preferential enrichment of pathways associated with metabolic reprogramming and stress responses, including hypoxia, fatty acid metabolism, glycolysis, OXPHOS, and ROS signalling, distinguishing them from other TME populations. Given the distinct transcriptional adaptations observed in BM tumor cells, we sought to uncover the transcription factors (TF) driving these upregulated genes and pathways. We applied SCENIC to over 106,149 single tumor cells across KT, ECM, and BM samples, identifying a set of TF regulons selectively activated in BM tumor cells (**Supplementary Fig. S2G**). FLI1, IKZF1, and JUN were among the highly activated regulons in the BM, where they were associated with the regulation of immune-modulating and inflammatory pathways (**Fig. 2F, Supplementary Fig. S2H; Table S10**). Then, we leveraged spatial transcriptomics data to spatially validate the expression of representative genes from key Hallmark pathways within the RCC brain metastasis TME **(Fig. 2G)**. Selected genes included *VEGFA*, and *INHA* for hypoxia; *GPX3*, and *SOD2* for reactive oxygen species; *MYC*, *RPL22*, and *RPL34* for MYC targets V1; *TNF*, *IL1B*, *CXCL1*, *CXCL2*, and *CCL20* for TNF-α/NF-κB signaling; and *VIM* and *VCAN* for the EMT pathway. This spatial analysis independently validated the transcriptional programs inferred from the snRNA-seq data (**Fig. 2G**). We then visualized the enrichment of these pathways in spatial transcriptomics, highlighting hypoxia (**Fig. 2H; Supplementary Fig. S2I**), reactive oxygen species **(Fig. 2I; Supplementary Fig. S2L)**, and EMT representative genes (**Supplementary Fig. S2J-K**). To explore cellular crosstalk within the RCC BM TME, we described top prioritized LRI between tumor cells and other cells. Notable interactions include SEMA4D-PLXNB1/2 (44–46) between T cells, NG cells, B cells and tumor cells; and NRXN2/3-FGFR4 (30,47) between NG cells and tumor cells, which have been implicated in promoting tumor growth and invasiveness **(Fig. 2J; Supplementary Fig. S2M**).

Given the enrichment of inflammation-related pathways in RCC BM tumor cells, we next focused on LRIs between tumor and myeloid cells in the BM TME to elucidate the role of myeloid cells in tumor progression and immune evasion. We identified interactions driving myeloid cell recruitment, polarization, and sustained immunosuppression, such as PROS1-MERTK (48–52) and C3-VSIG4(53–55) in BM TME (**Fig. 2K; Supplementary Fig. S2N**). Additionally, SAA1-TLR(56,57), MADCAM1-SIGLEC9(58,59) and SERPING1-LRP1(60) interactions suggest macrophage-mediated immune suppression. To validate the prioritized LRIs identified in snRNA-seq, we examined the expression of CD163, a marker of immunosuppressive myeloid cells, along with key ligand-receptor pairs, including SAA1-TLR2 with downstream target IL10 (61–65), MERTK with downstream target IL10(52), and SAA1-FPR1 with downstream targets S100A8 and S100A9 (66), all of which have been implicated in promoting an immunosuppressive niche **(Fig. 2L)**. We next used spatial transcriptomics to visualize these LRIs expression between neighboring tumor cells and myeloid populations **(Fig. 2M and Supplementary Fig. S2O; Supplementary Fig. S2P-Q**) confirming our findings. Neighborhood analysis of the spatial transcriptomics data revealed that myeloid cells were the predominant neighboring cell type surrounding 5,987 tumor cells within the brain metastasis microenvironment, comprising approximately 12% of all neighboring cells within a 85 µm neighborhood diameter (**Supplementary Fig. S2R**). Collectively, these findings demonstrate that RCC brain metastasis tumor cells, together with neighbouring TME populations, adopt a distinct inflammatory and metabolically reprogrammed state driven by both tumor-intrinsic transcriptional rewiring and coordinated interactions with the surrounding microenvironment.

### RCC Brain Metastasis Enrichment of Immunosuppressive Tumor Associate Macrophages

Among TME components, myeloid cells have been implicated as key modulators promoting immune evasion and tumor progression(67). To dissect the heterogeneity of tumor-associated myeloid populations across primary and metastatic sites, we performed unsupervised clustering and dimensionality reduction on 28,080 cells. Uniform manifold approximation and projection (UMAP) visualization of the myeloid population revealed a diverse landscape comprising monocytes, dendritic cells, and multiple macrophage subtypes (**Fig. 3A**). Marker gene expression defined the major subsets, including monocytes (*FCN1, VCAN, LYZ*), dendritic cells (DC) (*CLNK, IDO1*) with subcluster of proliferating DCs (*EZH2, POLQ, TOP2A, KNL1*), and tumor-associated macrophages (TAMs) (*CD163, MERTK*) with subclusters of CSF1R⁺ TAMs (*CSF1R, SELENOP, F13A, MS4A6A, MS4A4E*), SPP1⁺ high TAMs (*SPP1*, *PPARG*, *CCL3, CCL4, OLR1*), CXCL10⁺ TAMs (*CXCL10, GBP1, GBP5*) and proliferating TAMs (*EZH2, POLQ, TOP2A, KNL1*); (**Fig. 3B; Supplementary Fig. S3A-B; Table S11**). To profile immunoregulatory states within the myeloid compartment, we calculated M1 (pro-inflammatory) and M2 (anti-inflammatory) polarization activity scores using curated gene signatures (68). CXCL10⁺ TAMs exhibited the highest M1 polarization activity scores, consistent with a pro-inflammatory phenotype (**Fig. 3C, Supplementary Fig. S3C**), whereas SPP1⁺ and CSF1R⁺ TAMs demonstrated strong M2 polarization activity scores, reflecting an immunosuppressive, tumor-supportive state (**Fig. 3D, Supplementary Fig. S3C**). Consistent with these polarization profiles, Hallmark pathway analysis revealed that CXCL10⁺ TAMs were enriched for inflammatory programs, including the interferon alpha response, interferon gamma response, IL6-JAK-STAT3 signalling, and inflammatory response pathways (**Supplementary Fig. S3C**). In contrast, SPP1⁺ and CSF1R⁺ high TAMs were enriched for M2-related pathways, further reinforcing their roles in immunosuppressive remodelling of the TME. Moreover, proliferating myeloid and dendritic cell subsets demonstrated strong enrichment for E2F targets and mitotic spindle pathways, indicative of active cycling and expansion.

**Figure 3.**
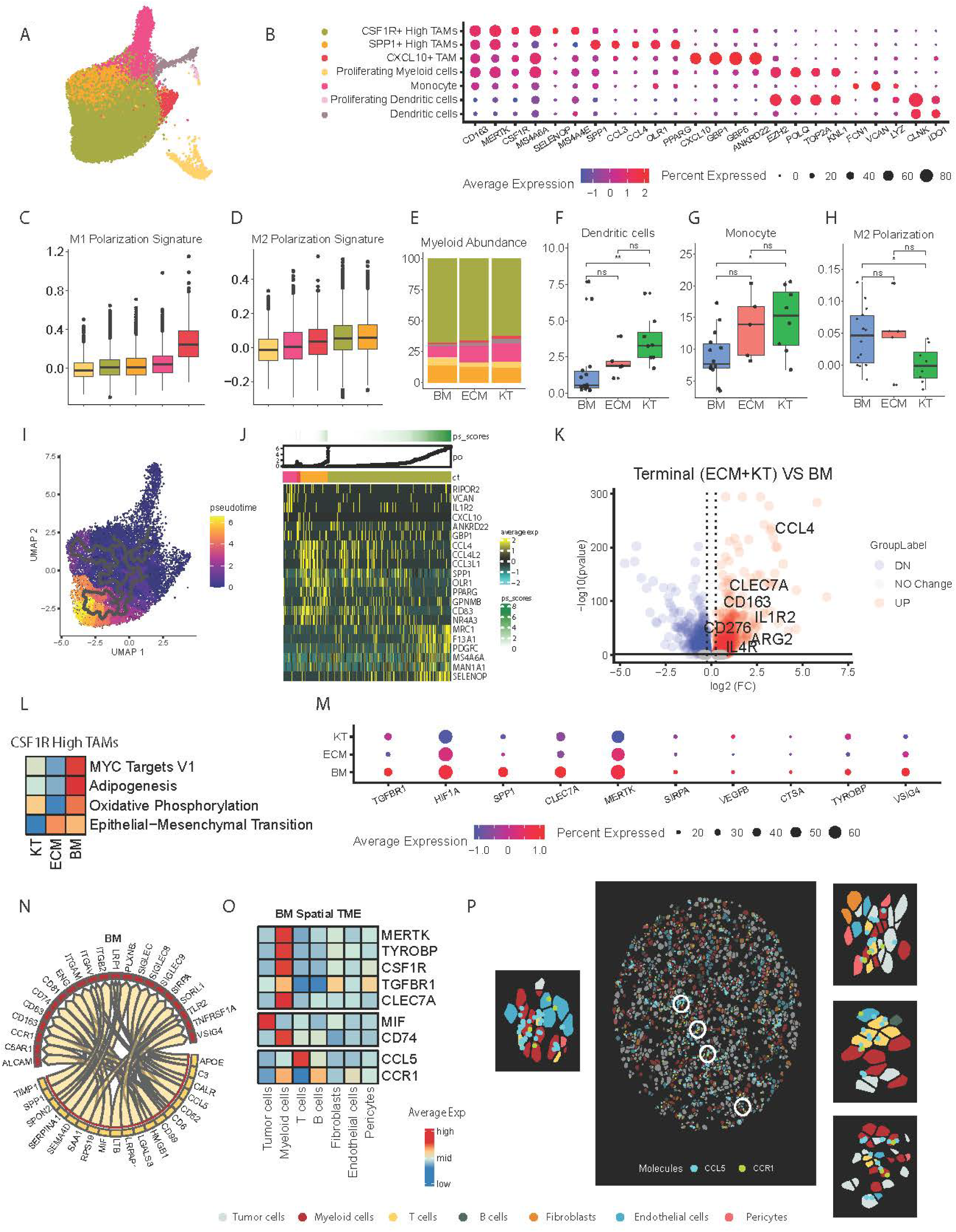
RCC Brain Metastasis Enrichment of Immunosuppressive Tumor Associate Macrophages. **(A)** UMAP representation of 28,080 myeloid cells across brain metastases (BM), extracranial metastases (ECM), and kidney tumors (KT), identifying 7 major subsets: monocytes, CXCL10⁺ TAMs, SPP1⁺ high TAMs, CSF1R⁺ high TAMs, proliferating myeloid cells, proliferating dendritic cells, and dendritic cells. **(B)** Dot plot displaying scaled average expression (color scale) and percent-expressing cells (dot size) of marker genes distinguishing the 7 annotated myeloid cell subtypes across all tumor compartments. **(C-D)** Boxplot showing the distribution of M1 polarization signature and M2 polarization signature activity scores across macrophages (CSF1R+ High TAM, SPP1+ High TAM, monocytes, proliferating myeloid cells, CXCL10+ High TAMs). **(E)** Stacked bar plot showing relative myeloid subtype proportions across all compartments (KT, ECM, and BM). **(F-G)** Boxplot displaying the relative abundance of dendritic cells and monocyte cells respectively across the 3 anatomical compartments BM, KT, and ECM. Statistical significance was assessed using a two-sided Wilcoxon rank-sum test. Boxplots depict the median (canter line), interquartile range (box limits), and whiskers extending up to 1.5x the interquartile range. **(H)** Boxplot displaying the M2 polarization activity scores across 3 anatomical compartments BM, KT, and ECM. Statistical significance was assessed using a two-sided Wilcoxon rank-sum test. Boxplots depict the median (canter line), interquartile range (box limits), and whiskers extending up to 1.5x the interquartile range. **(I)** Monocle-based pseudo time trajectory of myeloid cells, rooted in monocytes, revealing progressive macrophage differentiation toward the immunosuppressive phenotype (CSF1R High TAMs) representing the terminally differentiated phenotype in the brain TME. **(J)** Heatmap depicting gene expression dynamics across the pseudo time trajectory of myeloid cells within the RCC BM microenvironment. Distinct transcriptional modules are activated at different stages of microglial development, reflecting progressive cellular adaptation to the brain niche. **(K)** Volcano plot displaying differential gene expression between CSF1R High TAMs from BM VS KT and ECM, highlighting genes significantly upregulated or downregulated in the BM compartment. Statistical significance was determined using a two-sided Wilcoxon rank-sum test. **(L)** Heatmap displaying AddModuleScore-derived activity scores for statistically significant Hallmark pathways enriched from the FSGEA Enrichment analysis in CSF1R⁺ high TAMs across KT, ECM, and BM, highlighting compartment-specific enrichment of metabolic rewiring **(M)** Dot plot showing average expression (colour scale) and percent-expressing cells (dot size) of selected immunoregulatory genes across all TAMs populations from KT, ECM, and BM. **(N)** Circos plot showing the top prioritized ligand–receptor interactions between T cells and myeloid cells in the brain metastasis niche. The plot highlights directional signalling from T cells (senders) to myeloid cells (receivers), revealing compartment-specific communication axes that regulate myeloid cells activity within the brain TME. **(O)** Heatmap showing pseudobulk mean expression levels of well-studied, targetable immunoregulatory molecules across major cell types within the BM TME, derived from spatial transcriptomic data. **(P)** Spatial fields of view (FOVs) showing colocalization of targetable CCR1+ Myeloid cells nearby the CCL5+ T cells. Highlighting the close proximity between the 2 cell types within the BM TME.

Following these observations, we next quantified the abundance of myeloid subclusters across RCC sample sites (**Fig. 3E; Tablse S12**). SPP1⁺ and CSF1R⁺ immunosuppressive TAMs were consistently enriched across all compartments, whereas monocytes (P < 0.05) and antigen-presenting dendritic cells (P < 0.005) were significantly underrepresented in BM compared to KT (**Fig. 3F-G**). To further evaluate the immunosuppressive state, we calculated the M2 polarization signature across all TAMs. Although no significant differences were observed between BM and ECM, BM TAMs exhibited significantly higher M2 scores compared to KT (*P* < 0.05), reinforcing the distinct immunosuppressive nature of the BM TME (**Fig. 3H**). To explore how myeloid phenotypes evolve within distinct TMEs, pseudotime trajectory analysis revealed that, across all compartments, myeloid lineages progressed from monocytes toward a common terminal state predominantly composed of CSF1R⁺ high TAMs (**Fig. 3I-J, Supplementary Fig. S3D-G**), suggesting a conserved differentiation program across different compartments. These terminal macrophages consistently exhibited high expression of *MRC1*, *F13A1*, and *SELENOP*, markers associated with M2-like, anti-inflammatory phenotypes(69–73). To elucidate compartment-specific transcriptional adaptations underlying TAMs heterogeneity, we first performed DGEA comparing CSF1R⁺ high TAMs in the BM to their counterparts in KT and ECM. CSF1R⁺ high TAMs in the BM exhibited elevated expression of genes associated with alternative (M2-like) macrophage polarization, including (*ARG2, CD163, IL1R2, IL4R, CCL4, CLEC7A, and CD276*) **(Fig. 3K; Table S13)**, widely reported in the literature to reflect an anti-inflammatory M2 phenotype and correlated with poor clinical outcomes(74–78).

Pathway analysis revealed significant enrichment of multiple metabolic and immunosuppressive pathways. These included MYC_Targets V1 (*HSP90AB1, LDHA, GLO1, PPIA*) (79–81), OXPHOS *(CYC1, NDUFA1, ATP5PD, NDUFA2, ATP5F1B*)(82–84), Adipogenesis (*LPL, GPX3, GPX4*)(85), and EMT (*SERPINE1, SPP1, LOX, LGALS1*) (**Fig. 3L; Supplementary Fig. S3H-I; Table S14**). Pathway-level enrichment revealed a coordinated upregulation of genes linked to metabolic plasticity, immunoregulation, and stromal remodeling, defining a metabolically reprogrammed and immunosuppressive M2-like TAM phenotype within the BM compartment. (**Fig. 3K-L**). Based on M2-like TAM phenotype, we sought to identify immunoregulatory genes expressed in TAMs with therapeutic targetability. We identified VSIG4, MERTK, TYROBP, HIF1A, TGFBR1 are highly expressed therapeutic targets on BM TAMs including CSF1R+ M2 type TAMs (**Fig. 3M; Supplementary J)**. To further elucidate immunomodulatory crosstalk between the T cells and the myeloid population within the brain, we examined the prioritized LRIs (**Fig. 3N; Supplementary Fig. 3K**). Several immunosuppressive interactions were consistently observed across T cells and myeloid cells, including MIF-CD74, SAA1-TLR2, CCL5-CCR1 (86–88), and C3-VSIG4. Next, we validated therapeutic targets and key LRIs in the spatial transcriptomics. Within the BM TME, myeloid populations uniquely expressed MERTK, TYROBP, CSF1R, and TGFBR1 **(Fig. 3O; Supplementary Fig. S3L-M)**. Spatial mapping further confirmed the localization of key CCL5-CCR1 interaction between T cells and myeloid cells, important in recruitment of myeloid derived suppressor cells (86–88) **(Fig. 3O-P, Supplementary Fig. S3N-O**). These findings underscore that the RCC BM niche is defined by immunosuppressive, metabolically active TAM subsets whose transcriptional programs and LRIs provide targets for therapeutic intervention.

### Metabolic and Transcriptional Reprogramming of CD8⁺ T Cells in the RCC Brain Metastasis TME

Among the immune cells orchestrating antitumor immunity, CD8⁺ T cells serve as primary effectors of cytotoxic responses. However, in metastatic RCC, CD8⁺ T cells frequently exhibit profound dysfunction, limiting therapeutic efficacy (89). This impairment arises not only from chronic antigen stimulation and immunosuppressive signalling but also from metabolic stressors, including hypoxia, nutrient depletion, and lipid accumulation within the TME (90). To systematically define T cell subtypes across primary and metastatic sites, we profiled 20,377 intratumoral T cells across BM (*n* = 9,729), ECM (*n* = 3,049), and KT (*n* = 7,599). Unsupervised clustering was not able to transcriptionally distinct CD4⁺ and CD8⁺ T cell subpopulations given the limited expression of *CD4, CD8A CD8B* gene in snRNAseq data (**Supplementary Fig. 4B**). However, by using other canonical markers we defined seven T cell subsets: naive/memory T cells (*IL7R, BACH2, LEF1, TCF7, SELL*), NKT/γδ-like T cells (*CD247, GNLY, NCAM1, TRDC, FCG3A*), early activated T cells (*AOAH, GZMK, GZMA, GZMH, CCL5*), regulatory T (Treg) cells (*CTLA4, FOXP3, IL2RA*), proliferating cytotoxic T lymphocytes (CTLs) (*MKI67, TOP2A, EZH2*), and two exhausted CTLs clusters, named as 4-1BB-high CTL and 4-1BB-low CTL, expressing varying degree of exhaustion-related genes, *TOX*, *MIR155HG, TNFRSF9* (**Fig. 4A-B; Supplementary Fig. 4A-B; Table S15**) (91). To validate these annotations functionally, we applied curated gene signatures, calculating per single nucleus the activity of T cell exhaustion, anergy, CD8 activation, cytotoxicity, and related immune states (**Fig. 4C**) (92). 4-1BB-high CTLs exhibited the highest exhaustion and anergy levels, consistent with terminally dysfunctional phenotypes. The alignment of gene signatures with cluster identity confirms our transcriptomic-based annotations. Next, we quantified the abundance of these T cell subsets across sample sites (**Fig. 4D; Table S16)**. This analysis revealed less proliferating CTLs within BM compared to KT (*P* < 0.05), suggesting impaired proliferation and expansion of activated CTLs (**Fig. 4E; Table S17**). To further dissect T cell evolution across compartments, we performed trajectory analysis, initiating pseudotime trajectories from naive/memory T cells. In RCC BM, CD8⁺ T cells predominantly progressed toward a terminally exhausted 4-1BB-high CTL, marked by upregulation of *TOX, TNFRSF9*, and *MIR155HG* (**Fig. 4F-G**). In contrast, T cells from KT and ECM terminally differentiated into proliferating CTLs, characterized by enrichment of cell cycle-associated genes (*MKI67, TOP2A*) **(Supplementary Fig.S4C-I)**. These findings indicate that the brain TME T cells Favor exhausted states as terminal differentiation and have limited proliferation ability, potentially limiting effective antitumor immunity. To understand the underlying mechanisms of BM TME T cells adaptations and exhaustion, we next explored the transcriptional profile distinguishing 4-1BB-high CTL across compartments. We performed DGEA comparing BM-derived 4-1BB-high CTL to their counterparts in ECM and KT, followed by GSEA to contextualize the differentially expressed genes within pathway-level frameworks. This analysis revealed upregulation of multiple metabolic and stress-response pathways in BM-derived 4-1BB-high CTL, including MYC_Targets V1 (*LDHA*), Oxidative Phosphorylation (OXPHOS) (*COX7C*, *ATP5F1E*, *VDAC1*)(93,94), mTORC1 signaling (*GAPDH*, *TPI1*, *DDIT4*, *LDHA*), Glycolysis (*SLC16A3*, *LDHA*, *TPI1*)(95), and Hypoxia (*SERPINE1*, *TPI1*, *LDHA*)(96), indicating a shift toward a metabolically reprogrammed exhaustion state within the BM tumor microenvironment (**Fig. 4H-I; Supplementary Fig. S4J-K; Table S18-19**). In contrast, 4-1BB-high CTL from KT and ECM compartments exhibited enrichment for the Mitotic Spindle Hallmark pathway (*TLK1, FLNB, ARHGAP10*), reflecting a proliferative transcriptional program consistent with active cell cycling and clonal expansion (**Supplementary Fig. S4J-K**). Building on this observation, we next assessed all metabolic pathways activity across T cell populations from the three compartments using Hallmark gene sets. This analysis revealed that 4-1BB-high CTL were distinguished by elevated fatty acid metabolism, cholesterol accumulation (90), and oxidative phosphorylation (OXPHOS), indicative of a highly metabolically reprogramming association with exhaustion phenotype **(Fig. 4J)**(94). To investigate upstream regulators driving this metabolic and functional rewiring in BM TME, we applied SCENIC analysis to the entire T cells population across compartments. Brain-infiltrating T cells exhibited selective activation of transcriptional profiles linked to exhaustion and lipid stress, notably upregulation of XBP1 regulon, which is known as a key mediator of the unfolded protein response, endoplasmic reticulum stress, and lipid metabolism (**Fig. 4K**)(97–99). Given the clinical significance of immunoregulatory surface receptors, we next evaluated the expression of inhibitory and stimulatory checkpoints across all T cell populations (**Fig. 4L-M**) and specifically within the 4-1BB-high CTL subset (**Supplementary Fig. S4L-M**). BM-infiltrating T cells exhibited elevated expression of inhibitory receptors and reduced expression of co-stimulatory molecules. A similar pattern was seen in 4-1BB-high CTL, consistent with a dysfunctional and exhausted phenotype. To elucidate the interactions between the myeloid cells, tumor cells, and T cells in the BM, we identified LGALS3–LAG3 and CD86–CTLA4 as the top BM prioritized LRIs, known as well-established mediators of T cell dysfunction and immune escape within the TME(100–102) (**Fig. 4N; Supplementary Fig. S4N-P**). Finally, we validated spatial transcriptomics showed the spatial distribution of these immunoregulatory circuits within the TME. Receptors such as CTLA4 and LAG3 were predominantly expressed by T cells, while their cognate ligands, including LGALS3, and CD86, were enriched in tumor cells and myeloid cells, respectively (**Fig. 4.O-Q; Supplementary Fig. S4Q-R)**. While the brain environment drives T cells toward an exhausted state, we also observed that a higher proportion of TCF7⁺-naive/memory T cells in the BM was associated with better survival (hazard ratio [HR] = 0.16, 95% confidence interval [CI]: 0.03–0.83, *P* = 0.029; **Fig. 4R**). Together, these findings reveal a compartment-specific program of T cell exhaustion and metabolic reprogramming driven by the microenvironment, identifying vulnerabilities, and immune checkpoints that could inform therapeutic targeting strategies in RCC BM.

**Figure 4.**
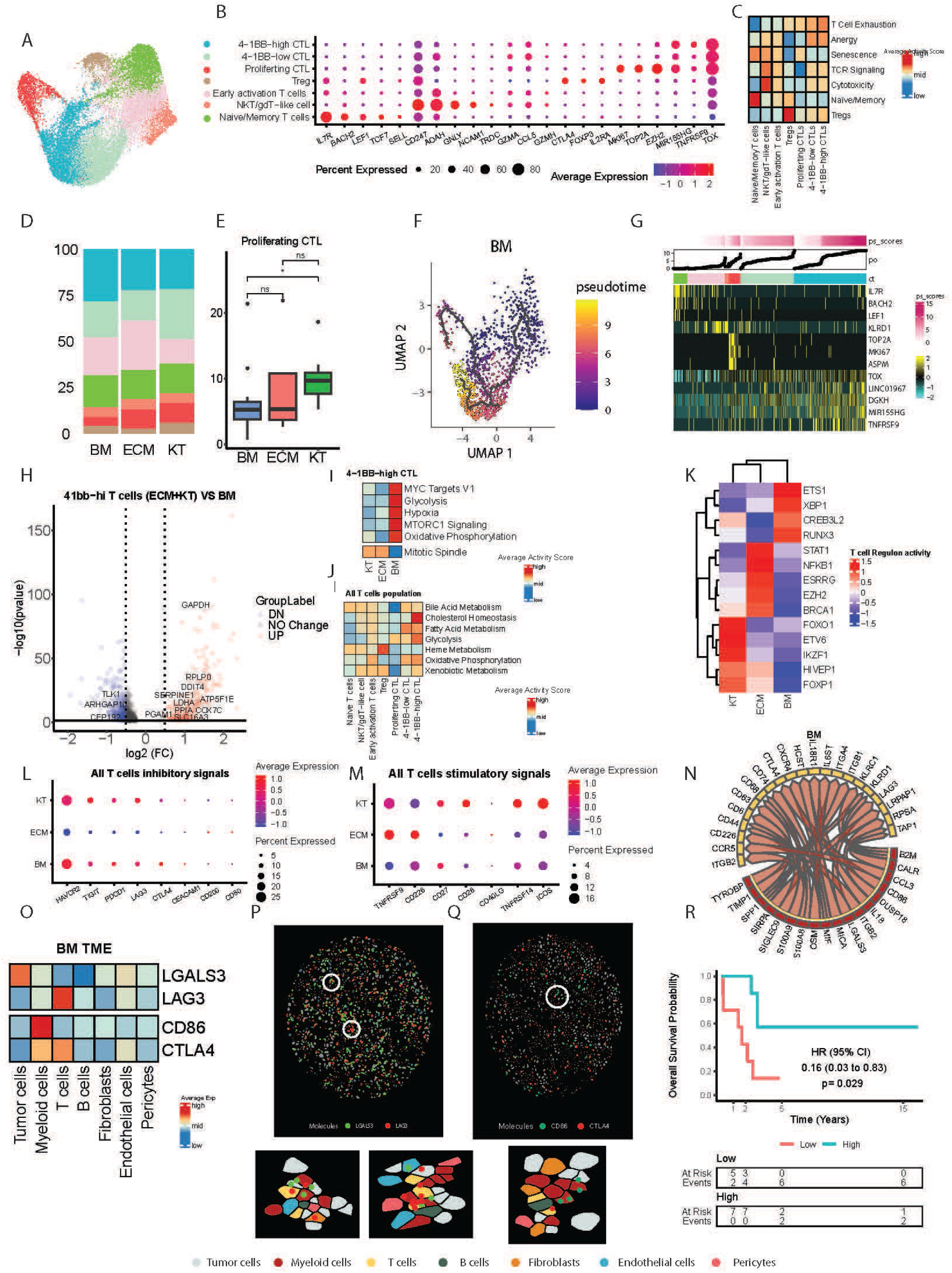
Metabolic and Transcriptional Reprogramming of CD8⁺ T Cells in the RCC Brain Metastasis TME. **(A)** UMAP representation of 20377 T cells across brain metastases (BM), extracranial metastases (ECM), and kidney tumors (KT), identifying 7 major subsets: Naive/Memory T cells,NKT/gdT−like cells,Early activation T cells,Tregs, Proliferting CTLs, 4−1BB−low CTLs, and 4−1BB−high CTLs. **(B)** Dot plot displaying scaled average expression (colour scale) and percent-expressing cells (dot size) of marker genes distinguishing the 7 annotated T cell subtypes across all tumor compartments. **(C)**Heatmap displaying AddModuleScore-derived mean activity scores for T cells specific signatures, confirming the T cell subtypes identified in the data. **(D)**Stacked bar plot showing relative T cells subtype proportions across all compartments (KT, ECM, and BM). **(E)** Boxplot displaying the relative abundance of Proliferating CTL across the 3 anatomical compartments BM, KT, and ECM. Statistical significance was assessed using a two-sided Wilcoxon rank-sum test. Boxplots depict the median (canter line), interquartile range (box limits), and whiskers extending up to 1.5x the interquartile range. **(F)** Monocle-based pseudo time trajectory of CD8+ T cells, rooted in Naive, revealing progressive T cells differentiation toward the 4−1BB−high CTLs phenotype, representing the terminally differentiated phenotype in the brain TME. **(G)** Heatmap depicting gene expression dynamics across the pseudo time trajectory of T cells within the RCC BM microenvironment. Distinct transcriptional modules are activated at different stages of T cells development. **(H)** Volcano plot displaying differential gene expression between 4−1BB−high CTLs from BM VS (KT & ECM), highlighting genes significantly upregulated or downregulated in the BM compartment. Statistical significance was determined using a two-sided Wilcoxon rank-sum test. **(I)** Heatmap displaying AddModuleScore-derived mean activity scores for statistically significant Hallmark pathways enriched from the FSGEA Enrichment analysis in 4−1BB−high CTLs across KT, ECM, and BM, highlighting compartment-specific enrichment of metabolic rewiring. **(J)** Heatmap displaying AddModuleScore-derived mean activity scores for metabolic Hallmark pathways in all T cells population across KT, ECM, and BM, highlighting the metabolic rewiring as the T cells develop from the naive towards the 4−1BB−high CTLs. **(K)** Heatmap displaying mean scaled regulon scores inferred by SCENIC in all T cells population across KT, ECM, and BM. **(L)** Dot plot displaying inhibitory signalling markers across T cell populations from the three compartments. Colour scale represents scaled average expression, and dot size reflects the percentage of expressing cells. **(M)** Dot plot displaying stimulatory signalling markers across T cell populations from the three compartments. Color scale represents scaled average expression, and dot size reflects the percentage of expressing cells. **(N)** Circos plot showing the top prioritized ligand–receptor interactions between T cells and myeloid cells in the brain metastasis niche. The plot highlights directional signalling from myeloid cells (senders) to T cells (receivers), revealing compartment-specific communication axes that regulate T cell activity within the brain TME. **(O)** Heatmap showing pseudobulk mean expression levels of well-studied, targetable immunoregulatory molecules associated with the T cells exhaustion across major cell types within the BM TME, derived from spatial transcriptomic data. **(P-Q)** Spatial fields of view (FOVs) showing colocalization of LGALS3+ Myeloid cells nearby the LAG3+ T cells, Moreover, CD86+ Myeloids neaby CTLA4+ T cells. Highlighting the close proximity between the 2 cell types within the BM TME. **(R)** Kaplan–Meier survival curves showing overall survival after craniotomy. Patients with high levels of naïve/memory T cells in brain metastases had significantly better outcomes, indicating these cells as a favourable prognostic marker.

### Brain metastasis microenvironment deficient of tertiary lymphoid of structures

To address the limited understanding of B cell function in metastatic sites, we profiled their phenotypes, distribution, and relationship to tertiary lymphoid structure (TLS) formation across different disease compartments. We performed unsupervised clustering and dimensionality reduction on 3,719 cells. UMAP visualization revealed 2 transcriptionally distinct B cell populations (**Fig. 5A**). To further delineate the transcriptional features distinguishing these clusters, we performed differential expression analysis. The first cluster was enriched for plasma and activated B cell markers, including *(CD38, XBP1, MZB1, PRDM1*), whereas the second cluster exhibited a naive/memory B cell transcriptional profile, characterized by elevated expression of (*MS4A1, BACH2, CD74, CD22)* (**Fig. 5B; Supplementary Fig. S5A; Table S20**). Beyond transcriptional differences, we next examined the numeric distribution of these B cell subsets across compartments. BM was enriched for activated/plasma B cells, while the ECM showed a relative enrichment of naive/memory B cells (**Fig. 5C; Table S21**). Activated/plasma B cells were significantly more abundant in the brain compared to ECM (*P* < 0.05) and KT microenvironments (*P* < 0.05) (**Fig. 5D; Table S22**). Naive/memory B cells were significantly enriched in the ECM compared to BM (*P* <0.05) (**Fig. 5E; Table S22**). Building on these transcriptional and numeric differences, we next investigated B cell LRIs in BM. We found that most of the BM TME cell types promote B cell differentiation toward an activated plasma cell state through engagement of the *SDC1* signalling axis (**Fig. 5F; Supplementary Fig. S5B**). Given the limited number of dendritic cells in BM, we sought to determine whether the activated/plasma cell differentiation occurs within mature tertiary lymphoid structures (TLS) or independently of TLS formation. As known, TLS, characterized by organized aggregates of B cells, T cells, and dendritic cells, is functionally significant and has been associated with improved responses to immune checkpoint blockade and favourable survival outcomes across multiple cancer types(103). In our selected 12 patient cohort with available whole-slide sections, we identified plasma cell aggregates in the absence of mature TLS structures within BM samples **(Fig.5G)**. In contrast, a representative lung metastasis sample contained several well-formed TLSs on a single slide (**Fig. 5H**). The absence of TLS formation in the brain may contribute to impaired antitumor immunity, resistance to immunotherapy, and the poor survival outcomes observed in patients with RCC and BM.

**Figure 5.**
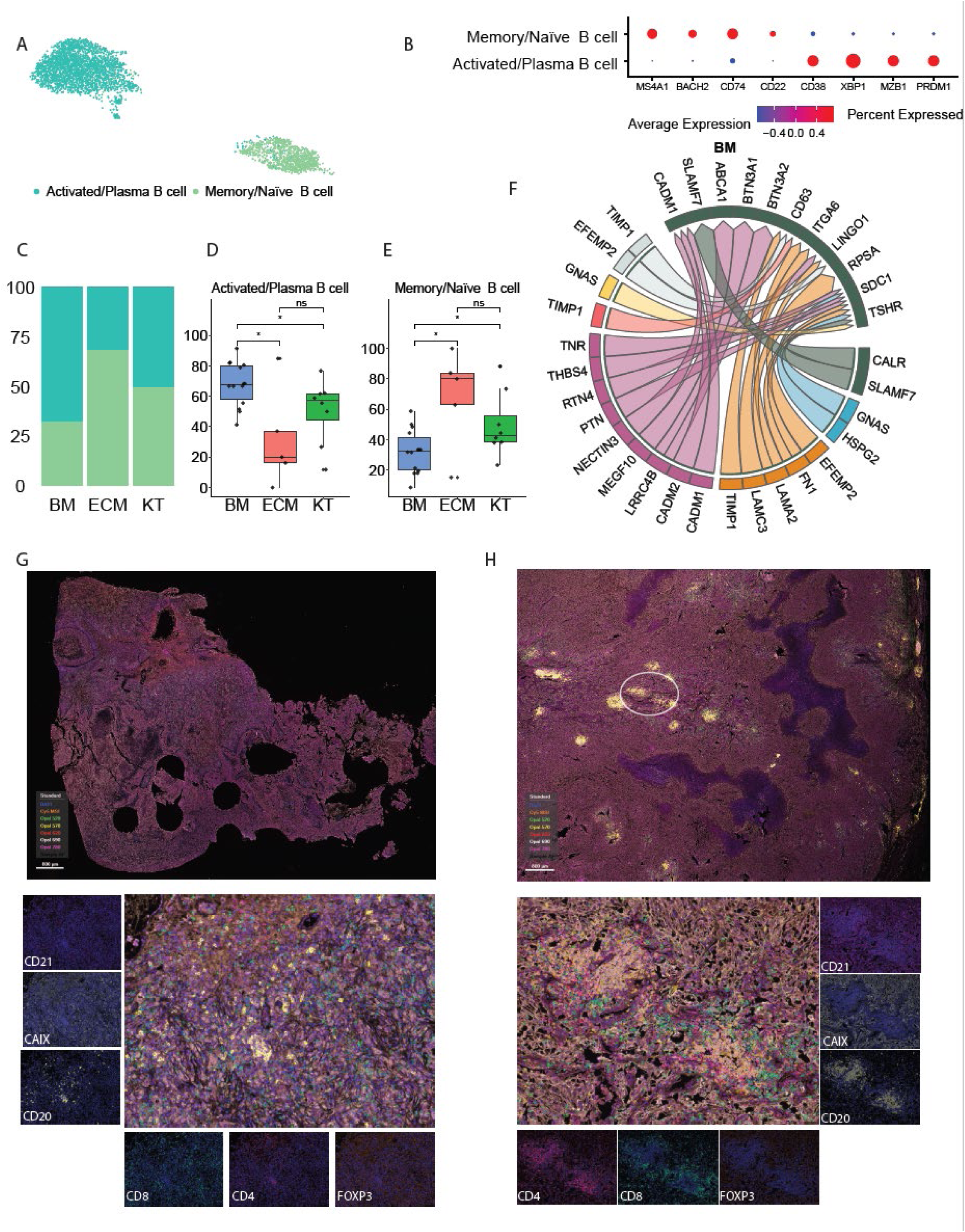
Brain metastasis microenvironment deficient of tertiary lymphoid of structures. **(A)** UMAP representation of 3,719 B cells across brain metastases (BM), extracranial metastases (ECM), and kidney tumors (KT), identifying 2 major subsets: activated/plasma cells and memory/naive B cells. **(B)** Dot plot displaying scaled average expression (colour scale) and percent-expressing cells (dot size) of marker genes distinguishing the 2 annotated B cell subtypes across all tumor compartments. **(C)** Stacked bar plot showing relative proportions of annotated B cell subsets across all compartments (KT, ECM, BM). **(D-E)** Boxplot displaying the relative abundance of activated/plasma B cells and the memory/naive B cells respectively across the 3 anatomical compartments BM, KT, and ECM. Statistical significance was assessed using a two-sided Wilcoxon rank-sum test. Boxplots depict the median (canter line), interquartile range (box limits), and whiskers extending up to 1.5x the interquartile range. **(F)** Circos plot showing prioritized ligand–receptor interactions mediating crosstalk between brain TME cell types and B cells in the RCC brain metastasis niche. The plot highlights directional signalling from TME cells (senders) to B cells (receivers), revealing compartment-specific communication axes that may regulate immune activity within the brain TME **(G)** Representative composite images showing the Opal multiplex immunofluorescence results employing the 7-plex RCC brain metastasis (BM) TLS panel with antibodies directed against CD4, CD8, FOXP3, CD20, CD21, CAIX, and DAPI. Whole-slide spatial plots display the distribution of CD20⁺ B cells and CD4⁺/CD8⁺ T cells, and their overlay with FOXP3⁺ regulatory T cells within the brain metastatic tumor microenvironment. **(H)** Representative composite images showing the Opal multiplex immunofluorescence results employing the 7-plex RCC lung metastasis (ECM) TLS panel with antibodies directed against CD4, CD8, FOXP3, CD20, CD21, CAIX, and DAPI. Whole-slide spatial plots display the distribution of CD20⁺ B cells and CD4⁺/CD8⁺ T cells, and their overlay with FOXP3⁺ regulatory T cells in the lung metastatic microenvironment.

### Stromal cells modulating brain metastasis immune microenvironment

Fibroblasts are key stromal cells involved in extracellular matrix organization and tissue remodelling. Beyond structural roles, they could modulate immune responses by secreting cytokines and chemokines. In cancer, fibroblasts could adopt immunosuppressive phenotypes that promote immune evasion and therapeutic resistance, particularly within metastatic niches(104). To explore fibroblast heterogeneity across tumor compartments, we performed unsupervised clustering and dimensionality reduction on 7,587 fibroblast cells. Of these, 2,870 fibroblasts were derived from the BM microenvironment, 1,853 from the KT TME, and 2,864 from ECM. Fibroblast clusters were annotated based on transcriptional programs previously defined in a pan-cancer fibroblast atlas(105), revealing 3 major subtypes: SPP1⁺ myofibroblasts (*SPP1, MET, TGFBI*), ACTA2⁺ myofibroblasts (*MYH11, ACTA2, TAGLN*), and adventitial fibroblasts (*MFAP5, PI16, CD34*) (**Fig. 6A-B**). Beyond transcriptional differences, we next examined the numeric distribution of these fibroblast subtypes across tumor compartments. No significant compartment-specific enrichment of fibroblast subpopulations was observed (**Fig. 6C; Table S23, Supplementary Fig. S6A-C; Table S24**). Given the transcriptional heterogeneity and distinct fibroblast subtypes, we performed trajectory analysis to see fibroblast differentiation. This analysis revealed that fibroblasts from all 3 compartments converged toward a terminal SPP1⁺ myofibroblast phenotype (**Fig. 6D; Supplementary Fig. S6D-E**). To further dissect this terminally differentiated population, we stratified SPP1⁺ myofibroblasts into 2 cohorts: those from BM and those from ECM and KT. Following DGEA and subsequent GSEA, we found that SPP1⁺ myofibroblasts in the BM TME were highly enriched for several Hallmark pathways, including MYC Targets V1(*HSP90AB1, LDHA, PGK1, RACK1*), Allograft Rejection (*HLA−A, HLA−E, HLA−DRA*), Interferon-α-Response (*IRF1, HLA−C, RIPK2*), and Hypoxia (*SLC2A1, SLC2A3, LDHA, TPI1, SERPINE1*). To obtain higher-resolution insights, we extended the analysis using Gene Ontology (GO) pathways. Similarly, this revealed that SPP1⁺ myofibroblasts in the BM TME exhibited strong enrichment of processes linked to inflammation, such as NF-κB activity (*IL18R1, HSPA1B, HSPA1A, PRKCQ*), positive regulation of the inflammatory response (*NFKBIZ, HLA−E, MAP3K8*), and cytoplasmic pattern-recognition receptor signalling (*TNFAIP3, RIPK2*). In contrast, SPP1⁺ myofibroblasts from ECM & KT TMEs were predominantly enriched for pathways associated with extracellular matrix biology, including ECM assembly (*COL1A2, COL3A1, ELN, THSD4*), ECM structural constituents (*MFAP5, COL15A1, COL12A1, COL1A2, COL3A1, COL8A1, LAMA3*), and ECM constituent conferring elasticity (*ELN, FBN2, FBLN5*) (**Fig. 6E-F; Table S25; Supplementary Fig. 6F-I; Table S26-27)**. These findings highlight distinct niche-driven inflammatory transcriptomic adaptation of SPP1⁺ myofibroblasts within the brain. To explore the communication of stromal cells to the surrounding immune cells within the brain,

**Figure 6.**
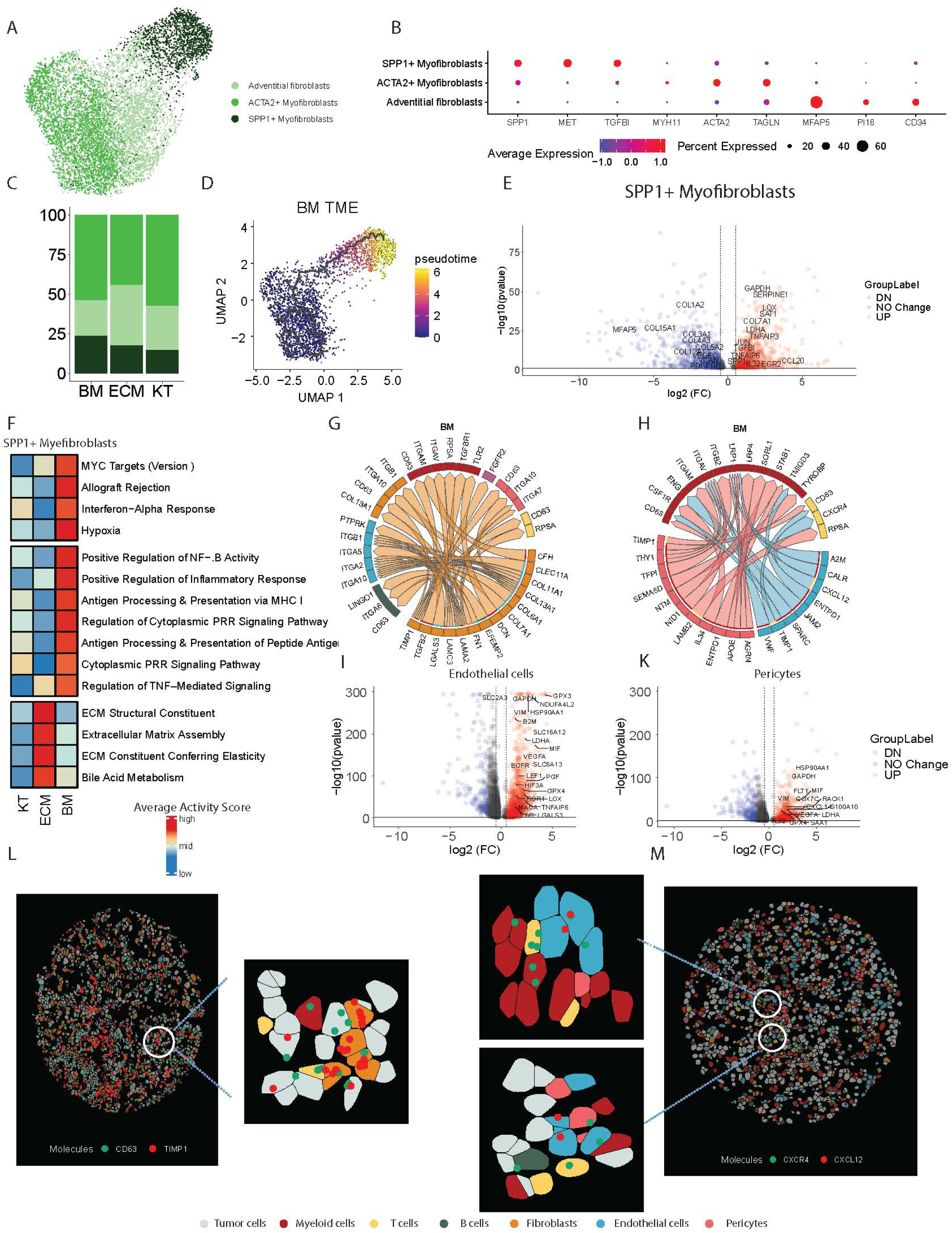
Stromal cells modulating brain metastasis immune microenvironment. **(A)** UMAP representation of 7587 fibroblast cells across brain metastases (BM), extracranial metastases (ECM), and kidney tumors (KT), identifying 3 major subsets: ACTA2+ myofibroblasts, adventitial fibroblasts, and SPP1+ myofibroblasts. **(B)** Dot plot displaying scaled average expression (colour scale) and percent-expressing cells (dot size) of marker genes distinguishing the 3 annotated fibroblast cell subtypes **(C)** Stacked bar plot showing relative proportions of annotated fibroblast cell subsets across all compartments (KT, ECM, BM). **(D)** Monocle-based pseudo time trajectory of fibroblast cells, rooted in ACTA2+ myofibroblasts, revealing progressive fibroblast differentiation across the BM TME. SPP1+ myofibroblasts represent the terminally differentiated phenotype in BM TME. **(E)** Volcano plot displaying differential gene expression between terminal differentiated SPP1+ myofibroblasts in BM vs. the terminal differentiated SPP1+ myofibroblasts in other sites (KT and ECM). Key upregulated and downregulated gene significance determined by two-sided Wilcoxon rank-sum test. **(F)** Pseudobulk heatmap showing scaled mean activity scores for Hallmark pathways and Gene Ontology pathways enriched from the FGSEA Enrichment analysis, the analysis shows the activity of these pathways per compartment. **(G)** Circos plot for the top prioritized ligand-receptor interactions mediating stromal-to-immune cell crosstalk within the brain TME of RCC metastases. The analysis highlights directional signalling from stromal cells (sender) to TME cell types (receiver), revealing compartment-specific communication axes that may shape immune cell behaviour in the BM niche. **(H)** Circos plot for the top prioritized ligand-receptor interactions mediating Endothelial and Pericytes-to-immune cell crosstalk within the brain TME of RCC metastases. The analysis highlights directional signalling from Endothelial and Pericytes (sender) to TME Immune cell types (receiver), revealing compartment-specific communication axes that may shape immune cell behaviour in the BM niche. **(I)** Volcano plot displaying differential gene expression in the endothelial cell population (14196 cells). Endothelial cells that come from the BM TME vs. those that come from the KT & ECM TMEs together. Significance is determined by two-sided Wilcoxon rank-sum test. **(K)** Volcano plot displaying differential gene expression pericytes population (5124 cells). Pericytes come from the BM TME vs. those from the KT & ECM TMEs together. Significance is determined by two-sided Wilcoxon rank-sum test. **(L)** Spatial fields of view (FOVs) showing colocalization of TIMP1+ Fibroblasts nearby the CD63+ T cells. Highlighting the close proximity between the 2 cell types within the BM TME. **(M)** Spatial fields of view (FOVs) showing colocalization of CXCL12+ Endothelial cells nearby the CXCR4+ T cells. Highlighting the close proximity between the 2 cell types within the BM TME.

We systematically inferred LRIs, identifying key signaling routes through which fibroblasts, pericytes, and endothelial cells influence T cell and myeloid cells. The analysis showed that the fibroblasts in BM are modulating the T cells and the myeloid cells through TIMP1-CD63 (106), and TGFB2-TGFBR1(107,108) interactions respectively. Additionally, the endothelial cells were also engaged in crosstalk with T cells via TIMP1-CD63(106) and CXC12-CXCR4(108–110) interactions. Furthermore, the pericytes engaged with the myeloid cells through IL34-CSF1R (111) (**Fig. 6G-H; Supplementary Fig. S6J-K**). After identifying stromal cell signals influencing immune populations, we next investigated transcriptional profiles of endothelial cells and pericytes across sample sites. We analysed a total of 5,794 endothelial cells derived from BM and 8,402 endothelial cells from KT and ECM. DGEA revealed that BM endothelial cells exhibited a distinct transcriptional profile compared to those from ECM and KT. Notably, endothelial cells within the BM TME exhibited strong enrichment for multiple immune-related Hallmark pathways, including TGF-beta signalling (*SERPINE1, ACVR1, RHOA*), TNF-alpha signalling (*TNFAIP8, TNFAIP6, IL6ST*) **(Supplementary Fig. 2E; Table S28)**. Similarly, pericytes from BM displayed a unique transcriptional program enriched for genes linked to inflammatory complement (*C1QA, SERPINA1, CLU*) and allograft rejection (*TLR2, HLA-A, B2M*) pathways. (**Fig. 6K; Supplementary Fig.S2C; Table S29**). Moreover, both endothelial cells and pericytes had enrichment of OXPHOS, lipid metabolism, ROS, hypoxia metabolic pathways consistent with previous findings **(Supplementary Fig. 2E)**. Collectively, these findings highlight that stromal cells within the brain undergo niche-specific transcriptional adaptations to modulate immune responses. We validated the spatial co-localization of stromal-T cell interactions, including CD63–TIMP1 and CXCL12–CXCR4 supporting their potential role in sustaining T cell dysfunction within the tumor microenvironment **(Fig.6L-M; Supplementary Fig.L-M).**

## DISCUSSION

Here, we provide a comprehensive characterization of the RCC BM TME through leveraging snRNA-seq and spatial transcriptomics technologies. Our study revealed BM TME-specific adaptation mechanisms across tumor, immune, and stromal cell types that contribute to immune evasion and therapeutic resistance. Analysis of LRIs in BM revealed a dominance of NG-to-NG signalling, consistent with the highly specialized cellular architecture of the brain (**Fig. 1G**). We found that RCC tumor cells in BM exhibited neuronal-like phenotypes, marked by expression of neural lineage and synaptic genes (**Supplementary Fig. S1L**), and engaged in active crosstalk with resident NG cells (**Fig. 1H; Supplementary Fig. S1K**). These findings reflect a broader phenomenon of neuronal mimicry, previously documented in other cancers, wherein metastatic cells adopt neuronal programs to enhance their survival and integration within the central nervous system (CNS) microenvironment(19,112–114). LRIs further supported this mimicry, identifying tumor-NG interactions involving classical pre- and post-synaptic molecules such as NLGNs-NRXN (**Fig. 1H; Supplementary Fig. S1K**). Collectively, These finding build the narrative of neuronal mimicry seen is other solid tumors to include RCC, showing that RCC can also exploit neural phenotype for metastatic adaptation and progression (114). In this context, neurons within the BM niche may further promote tumor aggressiveness through the secretion of metabolites that provide bioenergetic support and modulate signalling pathways, thereby reinforcing tumor growth and adaptation to the neural microenvironment. Importantly, such tumor –neuron interactions are also likely to influence the efficacy of immunotherapy, as metabolite-driven adaptations can impair T-cell function and contribute to immune evasion within the BM (115). We also identified semaphorin-mediated signalling, through SEMA4D-PLXNB2 and SEMA4D-MET interactions, as a dominant axis between NG and tumor cells. These pathways are well-characterized drivers of tumor proliferation, invasion, and immune modulation(116,117). In addition, we observed NRXN2-FGFR4 interactions in our LR analysis between NG and tumor cells. FGFR4 is a key oncogenic driver in RCC, implicated in promoting tumor cell proliferation, invasion, survival, and resistance to VEGF-targeted therapies (30,118). The engagement of NRXN2-FGFR4 and SEMA4D-MET by NG cell-derived ligands in the brain may represent a previously unrecognized mechanism through which NG cells contribute to tumor progression in BM.

Compared to KT and ECM, tumor cells in BM exhibited a distinct transcriptional profile that likely reflects adaptation to the brain microenvironment. These cells consistently upregulated pathways associated with metabolic stress and nutrient limitation, including glycolysis, oxidative phosphorylation, ROS, adipogenesis, mTORC1, and MYC target gene programs (**Fig. 2C; Supplementary Fig. S2B**). These tumor intrinsic transcriptional changes reflect a metabolically reprogrammed state and could affect the immune cells function in tumor microenvironment(119). Prior studies on BM have identified similar patterns of metabolic adaptation as essential mechanisms for tumor cells in the brain (120–124). The enrichment of MYC target gene signatures, together with MYC’s established role as a master regulator of metabolism, suggests that MYC may orchestrate the metabolic adaptations required for tumor survival and progression within the RCC BM microenvironment (125,126).

Consistent with the study by Hoover et al., which demonstrated that neurons can transfer mitochondria to cancer cells, we propose that neuroglial cells in RCC brain metastases may similarly promote metabolic reprogramming in tumor and surrounding cells, complementing the ligand–receptor interactions we observed. This reprogramming involves a metabolic shift toward oxidative phosphorylation(127). The transcriptional differences were further reflected at the regulatory level. BM tumor cells exhibited selective activation of transcription factor regulons such as JUN, IKZF1, and FLI1, which are associated with inflammatory signalling, immune modulation, and complement regulation. The tumor-intrinsic enrichment of these regulons likely contributes to reshaping and modulating the tumor microenvironment (TME), reinforcing pro-tumorigenic communication within the brain niche. (**Fig. 2F, Supplementary Fig, S2G-H**)(128,129).

Focusing on the incoming cues to tumor cells, we observed that not just only the neuroglial cells engaged with the tumor cells through SEMA4D/Plexin signalling but also T cells, and B cells. This LRIs are poised to enhance tumor cell adaptation and survival within the brain microenvironment. Consistent with its established roles in tumor growth, and immune modulation(46,117). Notably, therapeutic blockade of SEMA4D has been shown to augment immune cell infiltration and suppress tumor progression in preclinical models, highlighting its potential as a promising target in RCC-BM(117).

We also examined outgoing signals from tumor cells toward the immune microenvironment. Tumor-to-myeloid LRIs in BM prominently featured immunoregulatory axes such as PROS1-MERTK(51), C3-VSIG4(53), MADCAM1-SIGLEC 9(130), and SAA1-TLR2/FPR1(56,66) (**Fig. 2K**). These interactions contribute to tumor progression by fostering a tumor-supportive microenvironment and actively modulating the TME to Favor immune evasion and tumor growth, In RCC and other tumor types, MERTK(131) and VSIG4 (132) signalling have been associated with poor response to immune checkpoint blockade, and SAA1-TLR2(133) engagement has been linked to downstream activation of IL-10 and S100A8/9(134,135), further amplifying myeloid-driven suppression.

Having identified tumor-derived immunosuppressive signals directed at myeloid cells, we next considered how these interactions can shape the myeloid landscape across anatomical compartments. Although immunosuppressive TAMs were common across all sites, the brain TME was notably depleted of monocytes and dendritic cells, suggesting a selective exclusion or of antigen-presenting and pro-inflammatory subsets. Next, we examined the expression of several therapeutically targetable genes, including TGFBR1, HIF1A, SPP1, CLEC7A, MERTK, SIRPA, TYROBP, VEGFB, and VSIG4, within the TMEs. Notably, these targets exhibited markedly elevated expression in the BM TME. Among them, SPP1 and TGFBR1 were highly expressed in a large proportion of myeloid cells, specifically and uniquely enriched in the BM TME, highlighting their potential as brain metastasis–associated immunomodulatory targets (67).

Their compartment-specific enrichment suggests that these pathways may be selectively targetable in RCC BM. T cell-myeloid interactions in the brain were dominated by signals known to suppress antigen presentation, promote tolerance, and impair cytotoxic function. Axes such as MIF-CD74(136), SAA1-TLR2(56) and C3-VSIG4 (137) likely sustain a local immune state that is resistant to activation and poorly responsive to immunotherapy. To understand how immune dysfunction extends beyond myeloid cells, we next profiled T cells across compartments and identified distinct T cell subclusters with a gradual transition from naive to exhausted states. Trajectory inference of T cell populations using Monocle3 revealed a naive-to-effector continuum shaped by the TME, wherein BM favoured terminal differentiation into 4-1BB-high CTL exhibiting an exhausted phenotype. These findings underscore the pivotal role of the TME in dictating T cell fate decisions and functional states across metastatic compartments. Despite similar frequencies. 4-1BB-high CTLs in BM were enriched for metabolic pathways including glycolysis, and OxPhos, hypoxia, mTORC1 signalling, and MYC Targets V1, suggesting adaptation to the metabolically challenging brain niche. Such metabolic rewiring shared features with exhausted T cells across the three tumor microenvironments BM, ECM, and KT where enrichment of lipid metabolism and cholesterol homeostasis Hallmark pathways were observed. These metabolic adaptations collectively impair effector function and promote T-cell dysfunction and exhaustion within the tumor microenvironment **(Fig. 4J)** (90,138).

This compartmental imprinting was further reflected at the transcriptional regulator level. The XBP1 transcription factor, which is linked to metabolic stress and impaired antitumor immunity(90), was selectively active in brain-infiltrating T cells, while ECM showed EZH2, associated with T cell differentiation and proliferation was associated with the anti-tumor immunity(139). Inhibitory receptors including VSIR, HAVCR2, PDCD1, LAG3, TIGIT, and CTLA4 were highest in brain T cells, while co-stimulatory molecules were more prominent in ECM and KT **(Fig.4 L-M).** While T cells display signs of functional exhaustion, their interactions with immunosuppressive myeloid cells via LGALS3-LAG3, CD86-CTLA4, underscore the potential targets for combination therapies to disrupt this crosstalk **(Fig.4 N)**.

In addition to T and myeloid cell dysfunction, B cell composition in BM showed a site-specific shift. While KT and ECM contained naive and memory B cells, BM were dominated by plasma B cell population, in the absence of tertiary lymphoid structures. Given prior studies linking TLS to improved survival and immunotherapy response across multiple tumor types(103,140), the absence of organized lymphoid architecture in the brain may limit effective B cell-mediated immunity. Inferred SDC1 signalling from surrounding stromal and NG cells suggests that local crosstalk may promote B cell maturation and differentiation; in the absence of TLS, these plasma B cells may lack the functional context needed for sustained antitumor activity.

Following the compartment-specific immune changes, we assessed stromal cells across RCC sites and identified brain-specific features in fibroblasts, endothelial cells, and pericytes. SPP1⁺ myofibroblasts in BM displayed a distinct transcriptional profile marked by enrichment of inflammatory, immune modulatory pathways. This pattern mirrors the adaptation observed within the brain niche, suggesting a conserved response to local stressors. In contrast, SPP1⁺ myofibroblasts in ECM were enriched for extracellular matrix organization pathways. Fibroblasts showed extensive communication with surrounding stromal and immune cells. Notably, they engaged with microglia via TGFB2-TGFBR1(107,141) and with T cells via TIMP1-CD63(106), both of which are associated with immunosuppressive effects on their respective target populations. Endothelial cells in BM also contributed to immune exclusion through CXCL12-CXCR4 signalling, a well-established axis known to spatially restrict T cell infiltration and limit access to the tumor core(108–110). In addition, pericytes engaged myeloid cells through IL-34-CSF1R, a pathway linked to M2 polarization and reduced immunotherapy efficacy(111,142).

A potential limitation of this study is that most patients included in the cohort received high dose steroids as part of the standard care for managing the inflammation associated with the brain metastasis prior to the surgical resection. The use of steroid therapy might have influenced the transcriptomic profile of the tumor microenvironment by decreasing the inflammation. Despite this, our transcriptomic analysis revealed a highly inflamed tumor microenvironment within the brain metastases. This observation suggests that, even under steroid treatment, the inflammatory state persists, indicating that the anti-inflammatory effects of steroids may not be sufficient to substantially alter the overall immune landscape captured in our study. Another limitation stems from the current technical constraints of the CosMx SMI platform. Due to the limited availability of probes targeting neuroglial markers and rarity of cells, we were unable to comprehensively capture neuroglial cell populations within our spatial transcriptomic dataset. To overcome this limitation, we chose more standard protein level approach with NeuN antibody. Finally, although this represents the largest single cell RCC brain metastasis study to date, the cohort size (14 matched tumor samples) remains relatively limited for broad statistical generalization with potential confounding factors. Given the retrospective nature of the work, several potential confounding factors may have influenced the molecular and immune features observed. These include differences in patient gender, treatment history, time to brain metastasis development, duration of systemic treatment prior to sampling, tumor resection site, and underlying genomic alterations such as copy number variations (CNVs) that is known to alter TME (4). Such clinical and genomic heterogeneity could contribute to variability in immune composition and transcriptional states across samples, potentially driving distinct immune outcomes. Future studies incorporating larger, prospective, multi-institutional cohorts with harmonized clinical annotations and integrative genomic profiling will be essential to validate these findings and eliminate the influence of these confounding variables on the molecular programs shaping the brain metastasis microenvironment.

This study provides the first comprehensive spatial and transcriptomic landscape of RCC brain metastasis, enabling the identification of several targetable biomarkers within the metastatic microenvironment. Among these, NRXN4-FGFR4, and the SEMA4D–MET axis emerged as candidate therapeutic targets within tumor and stromal compartments, while MERTK-expressing tumor-associated macrophages (TAMs) were identified as critical mediators of immune modulation. Building on these findings, our integrative analyses have already informed the design of two RCC-focused clinical trials: one evaluating the combination of lenvatinib and pembrolizumab, aimed at targeting FGFR4-driven tumor signalling, and another testing zanzalintinib, designed to inhibit both tumor-intrinsic c-MET and macrophage-associated MERTK and CSF1R pathways. In addition to these translational advances, our study uncovered multiple metabolic and immunoregulatory biomarkers that warrant further mechanistic exploration. Future work should focus on elucidating how these metabolic adaptations contribute to immune evasion and therapy resistance, thereby providing a foundation for rational combination strategies that integrate metabolic, immune, and microenvironment-targeted interventions in RCC brain metastasis. In summary, this work presents the largest single-nucleus transcriptomic dataset of RCC brain metastasis to date, uncovering clinically actionable biomarkers and pathways that can inform the development of mechanism-based therapies.

## METHODS

### Sample Characteristics

We collected pathologically confirmed RCC samples from patients undergoing surgical resection at MD Anderson Cancer Center, with written informed consent and approval from the Institutional Review Board approved protocol. The study was conducted in accordance with ethical guidelines (Declaration of Helsinki). Specifically, 27 tumor samples from 14 patients including BM, ECM, and KT underwent snRNA-seq. and 12 BM samples from RCC patients were also profiled using CosMx SMI with a 900-plex targeted gene panel. De-identified data shared with Ohio State University investigators through Institutionally approved Data Transfer and Use Agreement.

### Nuclei Suspensions and Sorting

Flash-frozen tissue of approximately 5 mm × 5 mm in size was minced into fine pieces in cold DAPI/NST buffer + RNase inhibitor (NST-DAPI buffer with 0.1 U μl−1 RNase Inhibitor (NEB, M0314L) or 400 μl 1 M Tris-HCL pH 7.5, 80 μl 5 M NaCl, 120 μl 1 M MgCl2, 400 μl 10% NP-40, 39 ml DNase/RNase-free sterile H2O). Then, the minced tissue was transferred into 1.5-mm beads to isolate nuclei and run bead blaster at 4.0-M speed for 3 cycles, 20 s on, at 10-s intervals, and then transferred to ice for 5 min. The nuclei were then passed through a FAC filter (40 um) before FAC sorting on the BD Melody. Nuclei were sorted (200K-300K) with gating for both diploid and aneuploid peaks for all events above a noise threshold and a doublet threshold to remove clumps based on DAPI, forward and side scatter on the BD Melody System. Sorted cells were collected into 1.5-ml low-binding tubes inhibitor (coat using PBS+1% BSA) with 6-10ul RNase inhibitor.

### snRNA-seq Library Preparation

After sorting, the nuclei were centrifuged at 550g for 5 min at 4 °C. Supernatant was removed (leaving ∼20 ul), resuspend buffer was slowly added (PBS+BSA (1%) + 0.2 U/ul RNase Inhibitor) without resuspending the nuclei pellet. They were then incubated on ice for 5-10 min for buffer exchange. The nuclei were manually counted and diluted to within the recommended range (500-1000 nuclei/μl) at a volume to recover 7000-10000 nuclei for snRNA-seq. Single cell capture, barcoding, and library preparation were performed by following the 10× Genomics Single Cell Chromium 3’ protocols (CG000183, V3.1). Ten libraries were pooled to give a final concentration of 10 nM, and pooled samples were further qPCR for final concentration before submission for sequencing with the NovaSeq6000 sequencer using the S2 100 cycles flow cell with 28 cycles for read 1, 8 cycles for i7 index, and 91 cycles for read 2 through the ATGC core at MD Anderson.

### snRNA-seq Data Processing, Cell-type Annotation, Quality Control, and Data Visualization

Reads from single nuclei were demultiplexed and aligned using the GRCh38.p12 human genome reference using 10x Genomics Cell Ranger 3.1.0 pipeline (143–145). snRNA-seq data were processed using the Seurat package (v5.1.0). Unique molecular identifier (UMI) count matrices were normalized, scaled, and integrated across patient samples using the canonical correlation analysis to correct for batch effects. Principal component analysis (PCA) was performed, and the first 30 principal components were used for downstream clustering at a resolution of 0.5. Quality control was performed to retain nuclei with a minimum of 234 detected genes (nFeature_RNA ≥ 234) and 500 total counts (nCount_RNA ≥ 500). Nuclei with greater than 19.78% mitochondrial gene content were excluded.

### Cell Differentiation Trajectory Inference

To study transcriptional dynamics and infer cellular state transitions across key immune and stromal compartments, we applied Monocle3 (v1.3.7) for trajectory analysis. Trajectory graphs were learned using the learn_graph function, and cells were ordered along pseudotime using the order_cells function. Biologically relevant root populations were manually selected based on known cell biology for each lineage. Differential gene expression along pseudotime was assessed using Monocle3’s graph_test function with the "principal_graph" option. This approach enabled the identification of lineage-specific transcriptional rewiring.

### Prioritization of Immunoregulatory Ligand-receptor Interactions Using MultiNicheNetR

Cell-cell communication analysis was performed using the MultiNicheNetR framework to prioritize ligand-receptor interactions based on differential gene expression profiles. Differential expression analysis was conducted with the following thresholds: Log₂ fold-change (log2FC) > 0.5, *P*-value < 0.05, and minimum fraction of expressing cells > 5%. For each condition and comparison, the top 25 prioritized ligand-receptor interactions were selected to ensure consistency across all analyses and figures.

### Differential Gene Expression and Pathway Enrichment Analysis

Differential gene expression (DE) analysis was performed using the *FindAllMarkers*() function in the Seurat package (v5.1.0). Log2FC thresholds were standardized across analyses to a minimum log2FC of 0.25, and genes were considered significantly differentially expressed if they met an adjusted *P* value < 0.05. *P* value adjustment was performed using the Bonferroni correction method implemented within Seurat. All differential expression analyses were based on normalized and scaled RNA Assay.

### Gene Regulatory Network Analysis to Nominate Putative Metaprogram Regulators

Transcription factors (TFs) activity were characterized on a per-sample basis using the SCENIC R pipeline (v1.3.1), Spearman correlation networks between human transcription factors and potential target genes were computed using the runCorrelation() function. Gene regulatory networks were then inferred using GENIE3 (v1.26.0) with Random Forests (treeMethod = "RF"), specifying 50 trees per TF to balance computational efficiency and robustness. Subsequently, cis-regulatory motif analysis was performed using RcisTarget with human motif databases hg19-tss-centered-10kb-7species. Regulons were defined based on the top 5 regulators per target gene (coexMethod = "top5perTarget"). Regulon activity was quantified across single nuclei using AUCell, and mean area under the curve (AUC) scores were computed using AUCell (v.1.26.0).

### Nanostring CosMx platform specs

Spatial transcriptomic profiling was performed using the NanoString CosMx SMI platform with the 960-plex RNA panel, enabling high-resolution, single-molecule detection of 960 genes across tumor, immune, and stromal compartments. Standard CosMx protocols were followed for tissue preparation, imaging, and RNA detection at subcellular resolution, providing comprehensive spatial transcriptomic maps of RCC BM. Four markers DAPI, CD3, PanCK, and CD45 were used for cell segmentation. This process allowed for the identification of cell boundaries and the allocation of expressed transcripts at the cell level and the generation of a transcript-by-cell count matrix. Cells with a UMI count of fewer than 20 transcripts or 10 features were excluded from the analysis. After QC, a total of 13128 cells encompassing 12 FOVs remained. This corresponds to a median of 915 cells per FOV.

### Label Transfer Between snRNA-seq and CosMx Spatial Data

To annotate cell types within the CosMx SMI dataset, we applied a label transfer framework using the snRNA-seq dataset as the reference and the CosMx dataset as the query. Cell-type labels were transferred by first identifying anchors between datasets using the *FindTransferAnchors*() function in Seurat (v5.1.0), computed based on 30 principal components (dims = 30) from the PCAs embeddings of the snRNA-seq dataset. Following anchor identification, label transfer was performed, assigning to each CosMx cell. To validate the accuracy of label transfer, we conducted differential expression analysis between the predicted cell types within the CosMx dataset. The resulting expression profiles confirmed that each transferred cluster expressed canonical marker genes corresponding to their expected cell identity, supporting the fidelity of the label assignment.

### Cellular Neighbourhood Analysis

To characterize the spatial relationships among BM TME celltypes, we defined the neighborhoods using dbscan (v.1.2.0) for the RCC BM tumor cells in a 85-um radius. We then computed the cellular proportions of each cell type within these specified neighborhoods. Proportions of the neighboring cells are described in bar plots.

### Multispectral staining and imaging

Multispectral imaging using the Vectra Microwave treatment was applied to perform antigen retrieval, quench endogenous peroxidases, and remove antibodies from earlier staining procedures. The slides were stained with primary antibodies against NeuN, CAIX, Sox10, CD20, CD21, CD4, CD8, FOXP3 and tyramide signal amplification (TSA) dyes to generate Opal signal. The slides were scanned with the Vectra 3 image scanning system (Caliper Life Sciences), and signals were unmixed and reconstructed into a composite image with Vectra inForm software 2.4.8.

## Data and Code Availability

The raw sequencing data generated in this study will be deposited in the Gene Expression Omnibus (GEO) under accession number [xxx]. Processed snRNA-Seq datasets, metadata annotations, and supporting files will be made available through project-specific request to the corresponding author. The scripts used for the analysis are available at the following GitHub repository: https://github.com/hasanovlab (to be made public upon publication).

## DECLARATIONS

## Acknowledgements

E.H. was supported by the American Society of Clinical Oncology Conquer Cancer Foundation Young Investigator Award, the International Kidney Cancer Coalition Cecile and Ken Youner Scholarship, the Society for Immunotherapy of Cancer-NanoString Technologies Single Cell Biology Award, the Kidney Cancer Association Young Investigator Award, Gateway Cancer Research Foundation and the Pelotonia Institute for Immuno-Oncology Recruitment Startup Fund. Notably, the Society for Immunotherapy of Cancer-NanoString Technologies Single Cell Biology Award, the Kidney Cancer Association Young Investigator Award, and the Pelotonia Institute for Immuno-Oncology Startup Funds supported the correlative tissue sequencing and analyses conducted in this study. M.H. was supported by American Society of Clinical Oncology Conquer Cancer Foundation Young Investigator Award and start-up funds from The Ohio State University Comprehensive Cancer Center, which contributed to the analysis of this study.

We wish to acknowledge the Pathology Core within the Brain Tumor Research Program at MD Anderson Cancer Center supported through the SPORE in Brain Cancer award (P50CA127001) and the generous philanthropic contributions to the MD Anderson Cancer Neuroscience Program.

We also acknowledge CPRIT SINGLE CORE Facilities Grant (RP180684) for all single cell services and ATGC core grant CA016672(ATGC) and the NIH 1S10OD024977-01 for sequencing services in our manuscripts that include data generated by the facility.

## Authors’ Contributions

M.I.H.A. performed the primary and downstream analyses and wrote the first draft of the manuscript. Z.F.A. contributed to writing of the first draft. J.A.O.R. performed downstream analyses. A.K.C., J.L., and P.K.R. conducted initial data processing. A.K.C., and P.K.R. revised the first draft of the manuscript. D.J.H.S. revised the first draft of the manuscript. N.K. assisted in first draft preparation. T.N.A.L. and A.G.H. performed microscopy imaging studies. A.O.O., S.H., M.A.B., F.L., and J.T.H. contributed to tissue collection and study design, and L.M.N. contributed to tissue collection. T.M.T. and J.L. prepared the single-cell libraries. N.N. supervised single-cell library preparation and initial processing. M.H. partially funded the study and supervised downstream analyses. E.J. partially funded the study and supervised tissue collection. E.H. conceptualized and supervised the overall study, partially funded the study, supervised downstream analyses, and edited the first draft. All authors reviewed and approved the manuscript.

## REFERENCES

1. Siegel RL, Kratzer TB, Giaquinto AN, Sung H, Jemal A. Cancer statistics, 2025. CA Cancer J Clin. Wiley; 2025;75:10–45.

2. Motzer RJ, Jonasch E, Agarwal N, Alva A, Baine M, Beckermann K, et al. Kidney Cancer, version 3.2022, NCCN clinical practice Guidelines in oncology. J Natl Compr Canc Netw. Harborside Press, LLC; 2022;20:71–90.

3. Hasanov E, Yeboa DN, Tucker MD, Swanson TA, Beckham TH, Rini B, et al. An interdisciplinary consensus on the management of brain metastases in patients with renal cell carcinoma. CA Cancer J Clin. Wiley; 2022;72:454–89.

4. Fernández-Sanromán Á, Fendler A, Tan BJY, Cattin A-L, Spencer C, Thompson R, et al. Tracking nongenetic evolution from primary to metastatic ccRCC: TRACERx renal. Cancer Discov. 2025;OF1–23.

5. Gulati S, Barata PC, Elliott A, Bilen MA, Burgess EF, Choueiri TK, et al. Molecular analysis of primary and metastatic sites in patients with renal cell carcinoma. J Clin Invest [Internet]. American Society for Clinical Investigation; 2024;134. Available from: 10.1172/JCI176230

6. Li Y, Lih T-SM, Dhanasekaran SM, Mannan R, Chen L, Cieslik M, et al. Histopathologic and proteogenomic heterogeneity reveals features of clear cell renal cell carcinoma aggressiveness. Cancer Cell. Elsevier BV; 2023;41:139–163.e17.

7. Hasanov E, Gao J, Tannir NM. The immunotherapy revolution in kidney cancer treatment: Scientific rationale and first-generation results. Cancer J. Ovid Technologies (Wolters Kluwer Health); 2020;26:419–31.

8. Jonasch E, Hasanov E, Motzer RJ, Hariharan S, Choueiri TK, Huang B, et al. Evaluation of brain metastasis in JAVELIN Renal 101: Efficacy of avelumab + axitinib (A+Ax) versus sunitinib (S). J Clin Oncol. American Society of Clinical Oncology (ASCO); 2020;38:687–687.

9. Rathmell WK, Rumble RB, Van Veldhuizen PJ, Al-Ahmadie H, Emamekhoo H, Hauke RJ, et al. Management of metastatic clear cell renal cell carcinoma: ASCO guideline. J Clin Oncol. American Society of Clinical Oncology (ASCO); 2022;40:2957–95.

10. Flippot R, Dalban C, Laguerre B, Borchiellini D, Gravis G, Négrier S, et al. Safety and efficacy of nivolumab in brain metastases from renal cell carcinoma: Results of the GETUG-AFU 26 NIVOREN multicenter phase II study. J Clin Oncol. American Society of Clinical Oncology (ASCO); 2019;37:2008–16.

11. Hasanov E, Jonasch E. Management of brain metastases in metastatic renal cell carcinoma. Hematol Oncol Clin North Am. Elsevier; 2023;37:1005–14.

12. Takemura K, Lemelin A, Ernst MS, Wells JC, Saliby RM, El Zarif T, et al. Outcomes of patients with brain metastases from renal cell carcinoma receiving first-line therapies: Results from the International Metastatic Renal Cell Carcinoma Database Consortium. Eur Urol. Elsevier BV; 2024;86:488–92.

13. Andersen BM, Faust Akl C, Wheeler MA, Chiocca EA, Reardon DA, Quintana FJ. Glial and myeloid heterogeneity in the brain tumour microenvironment. Nat Rev Cancer. Springer Science and Business Media LLC; 2021;21:786–802.

14. Guan Z, Lan H, Cai X, Zhang Y, Liang A, Li J. Blood-brain barrier, cell junctions, and tumor microenvironment in brain metastases, the biological prospects and dilemma in therapies. Front Cell Dev Biol. Frontiers Media SA; 2021;9:722917.

15. Quail DF, Joyce JA. The microenvironmental landscape of brain tumors. Cancer Cell. 2017;31:326–41.

16. Fares J, Petrosyan E, Dmello C, Lukas RV, Stupp R, Lesniak MS. Rethinking metastatic brain cancer as a CNS disease. Lancet Oncol. Elsevier BV; 2025;26:e111–21.

17. de Carvalho Fraga CA, Tiburske L, Lucena da Silva GV, Simizo A, Cafundó de Morais MC, da Silva Fernandes Duarte AK, et al. Revealing shared molecular drivers of brain metastases from distinct primary tumors. Brain Res. Elsevier BV; 2025;1851:149456.

18. In GK, Ribeiro JR, Yin J, Xiu J, Bustos MA, Ito F, et al. Multi-omic profiling reveals discrepant immunogenic properties and a unique tumor microenvironment among melanoma brain metastases. NPJ Precis Oncol. Springer Science and Business Media LLC; 2023;7:120.

19. Tagore S, Caprio L, Amin AD, Bestak K, Luthria K, D’Souza E, et al. Single-cell and spatial genomic landscape of non-small cell lung cancer brain metastases. Nat Med. 2025;31:1351–63.

20. Hu C, Li T, Xu Y, Zhang X, Li F, Bai J, et al. CellMarker 2.0: an updated database of manually curated cell markers in human/mouse and web tools based on scRNA-seq data. Nucleic Acids Res. Oxford University Press (OUP); 2023;51:D870–6.

21. Thul PJ, Lindskog C. The human protein atlas: A spatial map of the human proteome. Protein Sci. 2018;27:233–44.

22. Browaeys R, Gilis J, Sang-Aram C, De Bleser P, Hoste L, Tavernier SJ, et al. MultiNicheNet: a flexible framework for differential cell-cell communication analysis from multi-sample multi-condition single-cell transcriptomics data [Internet]. bioRxiv. 2023. Available from: 10.1101/2023.06.13.544751

23. Palandri A, Salvador VR, Wojnacki J, Vivinetto AL, Schnaar RL, Lopez PHH. Myelin-associated glycoprotein modulates apoptosis of motoneurons during early postnatal development via NgR/p75(NTR) receptor-mediated activation of RhoA signaling pathways. Cell Death Dis. Springer Science and Business Media LLC; 2015;6:e1876.

24. Huang R, Yuan D-J, Li S, Liang X-S, Gao Y, Lan X-Y, et al. NCAM regulates temporal specification of neural progenitor cells via profilin2 during corticogenesis. J Cell Biol [Internet]. Rockefeller University Press; 2020;219. Available from: 10.1083/jcb.201902164

25. Aoto J, Martinelli DC, Malenka RC, Tabuchi K, Südhof TC. Presynaptic neurexin-3 alternative splicing trans-synaptically controls postsynaptic AMPA receptor trafficking. Cell. Elsevier BV; 2013;154:75–88.

26. Trotter JH, Wang CY, Zhou P, Nakahara G, Südhof TC. A combinatorial code of neurexin-3 alternative splicing controls inhibitory synapses via a trans-synaptic dystroglycan signaling loop. Nat Commun. 2023;14:1771.

27. Klimaschewski L, Claus P. Fibroblast growth factor signalling in the diseased nervous system. Mol Neurobiol. Springer Science and Business Media LLC; 2021;58:3884–902.

28. Pillai S, Netravali IA, Cariappa A, Mattoo H. Siglecs and immune regulation. Annu Rev Immunol. Annual Reviews; 2012;30:357–92.

29. Tamagnone L, Franzolin G. Targeting semaphorin 4D in cancer: A look from different perspectives. Cancer Res. American Association for Cancer Research (AACR); 2019. page 5146–8.

30. Narisawa T, Naito S, Ito H, Ichiyanagi O, Sakurai T, Kato T, et al. Fibroblast growth factor receptor type 4 as a potential therapeutic target in clear cell renal cell carcinoma. BMC Cancer. Springer Science and Business Media LLC; 2023;23:170.

31. Giordano S, Corso S, Conrotto P, Artigiani S, Gilestro G, Barberis D, et al. The semaphorin 4D receptor controls invasive growth by coupling with Met. Nat Cell Biol. Springer Science and Business Media LLC; 2002;4:720–4.

32. Tsetsenis T, Boucard AA, Araç D, Brunger AT, Südhof TC. Direct visualization of trans-synaptic neurexin-neuroligin interactions during synapse formation. J Neurosci. Society for Neuroscience; 2014;34:15083–96.

33. Cimadamore F, Amador-Arjona A, Chen C, Huang C-T, Terskikh AV. SOX2-LIN28/let-7 pathway regulates proliferation and neurogenesis in neural precursors. Proc Natl Acad Sci U S A. Proceedings of the National Academy of Sciences; 2013;110:E3017–26.

34. Han S, Okawa S, Wilkinson GA, Ghazale H, Adnani L, Dixit R, et al. Proneural genes define ground-state rules to regulate neurogenic patterning and cortical folding. Neuron. Elsevier BV; 2021;109:2847–2863.e11.

35. Wilkinson G, Dennis D, Schuurmans C. Proneural genes in neocortical development. Neuroscience. Elsevier BV; 2013;253:256–73.

36. Panagiotakos G, Pasca SP. A matter of space and time: Emerging roles of disease-associated proteins in neural development. Neuron. Elsevier BV; 2022;110:195–208.

37. Dhanasekaran R, Deutzmann A, Mahauad-Fernandez WD, Hansen AS, Gouw AM, Felsher DW. The MYC oncogene - the grand orchestrator of cancer growth and immune evasion. Nat Rev Clin Oncol. Springer Science and Business Media LLC; 2022;19:23–36.

38. Jin Y, Qiu J, Lu X, Li G. C-MYC inhibited ferroptosis and promoted immune evasion in ovarian cancer cells through NCOA4 mediated ferritin autophagy. Cells. MDPI AG; 2022;11:4127.

39. Wang Z, Goto Y, Allevato MM, Wu VH, Saddawi-Konefka R, Gilardi M, et al. Disruption of the HER3-PI3K-mTOR oncogenic signaling axis and PD-1 blockade as a multimodal precision immunotherapy in head and neck cancer. Nat Commun. Springer Science and Business Media LLC; 2021;12:2383.

40. Ma S, Zhao Y, Lee WC, Ong L-T, Lee PL, Jiang Z, et al. Hypoxia induces HIF1α-dependent epigenetic vulnerability in triple negative breast cancer to confer immune effector dysfunction and resistance to anti-PD-1 immunotherapy. Nat Commun. Springer Science and Business Media LLC; 2022;13:4118.

41. Ashton TM, McKenna WG, Kunz-Schughart LA, Higgins GS. Oxidative phosphorylation as an emerging target in cancer therapy. Clin Cancer Res. 2018;24:2482–90.

42. Li W, Chen S, Lu J, Mao W, Zheng S, Zhang M, et al. YY1 enhances HIF-1α stability in tumor-associated macrophages to suppress anti-tumor immunity of prostate cancer in mice. Nat Commun. Springer Science and Business Media LLC; 2025;16:6261.

43. Liu P, Sun S-J, Ai Y-J, Feng X, Zheng Y-M, Gao Y, et al. Elevated nuclear localization of glycolytic enzyme TPI1 promotes lung adenocarcinoma and enhances chemoresistance. Cell Death Dis. Springer Science and Business Media LLC; 2022;13:205.

44. Sierra JR, Corso S, Caione L, Cepero V, Conrotto P, Cignetti A, et al. Tumor angiogenesis and progression are enhanced by Sema4D produced by tumor-associated macrophages. J Exp Med. Rockefeller University Press; 2008;205:1673–85.

45. Franzolin G, Brundu S, Cojocaru CF, Curatolo A, Ponzo M, Mastrantonio R, et al. PlexinB1 inactivation reprograms immune cells in the tumor microenvironment, inhibiting breast cancer growth and metastatic dissemination. Cancer Immunol Res. American Association for Cancer Research (AACR); 2024;12:1286–301.

46. Evans EE, Paris M, Smith ES, Zauderer M. Immunomodulation of the tumor microenvironment by neutralization of Semaphorin 4D. Oncoimmunology. Informa UK Limited; 2015;4:e1054599.

47. Levine KM, Ding K, Chen L, Oesterreich S. FGFR4: A promising therapeutic target for breast cancer and other solid tumors. Pharmacol Ther. Elsevier BV; 2020;214:107590.

48. Ubil E, Caskey L, Holtzhausen A, Hunter D, Story C, Earp HS. Tumor-secreted Pros1 inhibits macrophage M1 polarization to reduce antitumor immune response. J Clin Invest. American Society for Clinical Investigation; 2018;128:2356–69.

49. Lahey KC, Gadiyar V, Hill A, Desind S, Wang Z, Davra V, et al. Mertk: An emerging target in cancer biology and immuno-oncology. Int Rev Cell Mol Biol. 2022;368:35–59.

50. Tanim KM, Holtzhausen A, Thapa A, Huelse JM, Graham DK, Earp HS. MERTK inhibition as a targeted novel cancer therapy. Int J Mol Sci. MDPI AG; 2024;25:7660.

51. Zhou Y, Fei M, Zhang G, Liang W-C, Lin W, Wu Y, et al. Blockade of the phagocytic receptor MerTK on tumor-associated macrophages enhances P2X7R-dependent STING activation by tumor-derived cGAMP. Immunity. Elsevier BV; 2020;52:357–373.e9.

52. Giroud P, Renaudineau S, Gudefin L, Calcei A, Menguy T, Rozan C, et al. Expression of TAM-R in human immune cells and unique regulatory function of MerTK in IL-10 production by tolerogenic DC. Front Immunol. Frontiers Media SA; 2020;11:564133.

53. Zheng X, Tong T, Duan L, Ma Y, Lan Y, Shao Y, et al. VSIG4 induces the immunosuppressive microenvironment by promoting the infiltration of M2 macrophage and Tregs in clear cell renal cell carcinoma. Int Immunopharmacol. Elsevier BV; 2024;142:113105.

54. Li J, Diao B, Guo S, Huang X, Yang C, Feng Z, et al. VSIG4 inhibits proinflammatory macrophage activation by reprogramming mitochondrial pyruvate metabolism. Nat Commun. 2017;8:1322.

55. Sazinsky S, Zafari M, Klebanov B, Ritter J, Nguyen PA, Phennicie RT, et al. Antibodies targeting human or mouse VSIG4 repolarize tumor-associated macrophages providing the potential of potent and specific clinical anti-tumor response induced across multiple cancer types. Int J Mol Sci. MDPI AG; 2024;25:6160.

56. Niu X, Yin L, Yang X, Yang Y, Gu Y, Sun Y, et al. Serum amyloid A 1 induces suppressive neutrophils through the Toll-like receptor 2-mediated signaling pathway to promote progression of breast cancer. Cancer Sci. Wiley; 2022;113:1140–53.

57. He M, Liu Y, Chen S, Deng H, Feng C, Qiao S, et al. Serum amyloid A promotes glycolysis of neutrophils during PD-1 blockade resistance in hepatocellular carcinoma. Nat Commun. Springer Science and Business Media LLC; 2024;15:1754.

58. Ibarlucea-Benitez I, Weitzenfeld P, Smith P, Ravetch JV. Siglecs-7/9 function as inhibitory immune checkpoints in vivo and can be targeted to enhance therapeutic antitumor immunity. Proc Natl Acad Sci U S A. Proceedings of the National Academy of Sciences; 2021;118:e2107424118.

59. van de Wall S, Santegoets KCM, van Houtum EJH, Büll C, Adema GJ. Sialoglycans and Siglecs can shape the tumor immune microenvironment. Trends Immunol. Elsevier BV; 2020;41:274–85.

60. Staudt ND, Jo M, Hu J, Bristow JM, Pizzo DP, Gaultier A, et al. Myeloid cell receptor LRP1/CD91 regulates monocyte recruitment and angiogenesis in tumors. Cancer Res. American Association for Cancer Research (AACR); 2013;73:3902–12.

61. Saraiva M, O’Garra A. The regulation of IL-10 production by immune cells. Nat Rev Immunol. Springer Science and Business Media LLC; 2010;10:170–81.

62. Kaynak A, Vallabhapurapu SD, Davis HW, Smith EP, Muller P, Vojtesek B, et al. TLR2-bound cancer-secreted Hsp70 induces MerTK-mediated immunosuppression and tumorigenesis in solid tumors. Cancers (Basel). MDPI AG; 2025;17:450.

63. Nguyen BN, Chávez-Arroyo A, Cheng MI, Krasilnikov M, Louie A, Portnoy DA. TLR2 and endosomal TLR-mediated secretion of IL-10 and immune suppression in response to phagosome-confined Listeria monocytogenes. PLoS Pathog. Public Library of Science (PLoS); 2020;16:e1008622.

64. Netea MG, Sutmuller R, Hermann C, Van der Graaf CAA, Van der Meer JWM, van Krieken JH, et al. Toll-like receptor 2 suppresses immunity against Candida albicans through induction of IL-10 and regulatory T cells. J Immunol. The American Association of Immunologists; 2004;172:3712–8.

65. Cheng N, He R, Tian J, Ye PP, Ye RD. Cutting edge: TLR2 is a functional receptor for acute-phase serum amyloid A. J Immunol. The American Association of Immunologists; 2008;181:22–6.

66. Wang Z, Wang Y, Yan Q, Cai C, Feng Y, Huang Q, et al. FPR1 signaling aberrantly regulates S100A8/A9 production by CD14+FCN1hi macrophages and aggravates pulmonary pathology in severe COVID-19. Commun Biol. Springer Science and Business Media LLC; 2024;7:1321.

67. Goswami S, Anandhan S, Raychaudhuri D, Sharma P. Myeloid cell-targeted therapies for solid tumours. Nat Rev Immunol. Springer Science and Business Media LLC; 2023;23:106–20.

68. Azizi E, Carr AJ, Plitas G, Cornish AE, Konopacki C, Prabhakaran S, et al. Single-cell map of diverse immune phenotypes in the breast tumor microenvironment. Cell. 2018;174:1293–1308.e36.

69. Song X, Zhu Y, Geng W, Jiao J, Liu H, Chen R, et al. Spatial and single-cell transcriptomics reveal cellular heterogeneity and a novel cancer-promoting Treg cell subset in human clear-cell renal cell carcinoma. J Immunother Cancer. BMJ; 2025;13:e010183.

70. Hughes R, Qian B-Z, Rowan C, Muthana M, Keklikoglou I, Olson OC, et al. Perivascular M2 macrophages stimulate tumor relapse after chemotherapy. Cancer Res. 2015;75:3479–91.

71. Opzoomer JW, Anstee JE, Dean I, Hill EJ, Bouybayoune I, Caron J. Macrophages orchestrate the expansion of a proangiogenic perivascular niche during cancer progression. Sci Adv. AAAS). American Association for the Advancement of Science; 2021;7.

72. Yang C, Qu J, Wu J, Cai S, Liu W, Deng Y, et al. Single-cell dissection reveals immunosuppressive F13A1+ macrophage as a hallmark for multiple primary lung cancers. Clin Transl Med. Wiley; 2024;14:e70091.

73. Du Y, Xia Y, Xu T, Hu H, He Y, Zhang M, et al. Selenoprotein o as a regulator of macrophage metabolism in selenium deficiency-induced lung inflammation. Int J Biol Macromol. Elsevier BV; 2024;281:136232.

74. Dowling JK, Afzal R, Gearing LJ, Cervantes-Silva MP, Annett S, Davis GM, et al. Mitochondrial arginase-2 is essential for IL-10 metabolic reprogramming of inflammatory macrophages. Nat Commun. Springer Science and Business Media LLC; 2021;12:1460.

75. Xia J, Zhang L, Peng X, Tu J, Li S, He X, et al. IL1R2 blockade alleviates immunosuppression and potentiates anti-PD-1 efficacy in triple-negative breast cancer. Cancer Res. American Association for Cancer Research (AACR); 2024;84:2282–96.

76. Zhang J, Dong Y, Yu S, Hu K, Zhang L, Xiong M, et al. IL-4/IL-4R axis signaling drives resistance to immunotherapy by inducing the upregulation of Fcγ receptor IIB in M2 macrophages. Cell Death Dis. Springer Science and Business Media LLC; 2024;15:500.

77. Wang J, Li X, Wang K, Li K, Gao Y, Xu J, et al. CLEC7A regulates M2 macrophages to suppress the immune microenvironment and implies poorer prognosis of glioma. Front Immunol. Frontiers Media SA; 2024;15:1361351.

78. Cheng M, Chen S, Li K, Wang G, Xiong G, Ling R, et al. CD276-dependent efferocytosis by tumor-associated macrophages promotes immune evasion in bladder cancer. Nat Commun. 2024;15:2818.

79. Pello OM, De Pizzol M, Mirolo M, Soucek L, Zammataro L, Amabile A, et al. Role of c-MYC in alternative activation of human macrophages and tumor-associated macrophage biology. Blood. American Society of Hematology; 2012;119:411–21.

80. Khan F, Lin Y, Ali H, Pang L, Dunterman M, Hsu W-H, et al. Lactate dehydrogenase A regulates tumor-macrophage symbiosis to promote glioblastoma progression. Nat Commun. Springer Science and Business Media LLC; 2024;15:1987.

81. Bae S, Park PSU, Lee Y, Mun SH, Giannopoulou E, Fujii T, et al. MYC-mediated early glycolysis negatively regulates proinflammatory responses by controlling IRF4 in inflammatory macrophages. Cell Rep. Elsevier BV; 2021;35:109264.

82. Wculek SK, Heras-Murillo I, Mastrangelo A, Mañanes D, Galán M, Miguel V, et al. Oxidative phosphorylation selectively orchestrates tissue macrophage homeostasis. Immunity. Elsevier BV; 2023;56:516–530.e9.

83. Wang F, Zhang S, Vuckovic I, Jeon R, Lerman A, Folmes CD, et al. Glycolytic stimulation is not a requirement for M2 macrophage differentiation. Cell Metab. Elsevier BV; 2018;28:463–475.e4.

84. Wang S, Liu G, Li Y, Pan Y. Metabolic reprogramming induces macrophage polarization in the tumor microenvironment. Front Immunol. Frontiers Media SA; 2022;13:840029.

85. Boufaied N, Chetta P, Hallal T, Cacciatore S, Lalli D, Luthold C, et al. Obesogenic high-fat diet and MYC cooperate to promote lactate accumulation and tumor microenvironment remodeling in prostate cancer. Cancer Res. American Association for Cancer Research (AACR); 2024;84:1834–55.

86. Li B-H, Garstka MA, Li Z-F. Chemokines and their receptors promoting the recruitment of myeloid-derived suppressor cells into the tumor. Mol Immunol. Elsevier BV; 2020;117:201–15.

87. Li B, Zhang S, Huang N, Chen H, Wang P, Yang J, et al. CCL9/CCR1 induces myeloid-derived suppressor cell recruitment to the spleen in a murine H22 orthotopic hepatoma model. Oncol Rep. Spandidos Publications; 2019;41:608–18.

88. Le K, Sun J, Ghaemmaghami J, Smith MR, Ip WKE, Phillips T, et al. Blockade of CCR1 induces a phenotypic shift in macrophages and triggers a favorable antilymphoma activity. Blood Adv. American Society of Hematology; 2023;7:3952–67.

89. Braun DA, Street K, Burke KP, Cookmeyer DL, Denize T, Pedersen CB, et al. Progressive immune dysfunction with advancing disease stage in renal cell carcinoma. Cancer Cell. Elsevier BV; 2021;39:632–648.e8.

90. Ma X, Bi E, Lu Y, Su P, Huang C, Liu L, et al. Cholesterol induces CD8+ T cell exhaustion in the tumor microenvironment. Cell Metab. Elsevier BV; 2019;30:143–156.e5.

91. Andreatta M, Corria-Osorio J, Müller S, Cubas R, Coukos G, Carmona SJ. Interpretation of T cell states from single-cell transcriptomics data using reference atlases. Nat Commun. Springer Science and Business Media LLC; 2021;12:2965.

92. Chu Y, Dai E, Li Y, Han G, Pei G, Ingram DR, et al. Pan-cancer T cell atlas links a cellular stress response state to immunotherapy resistance. Nat Med. 2023;29:1550–62.

93. Wu H, Zhao X, Hochrein SM, Eckstein M, Gubert GF, Knöpper K, et al. Mitochondrial dysfunction promotes the transition of precursor to terminally exhausted T cells through HIF-1α-mediated glycolytic reprogramming. Nat Commun. Springer Science and Business Media LLC; 2023;14:6858.

94. Li F, Liu H, Zhang D, Ma Y, Zhu B. Metabolic plasticity and regulation of T cell exhaustion. Immunology. Wiley; 2022;167:482–94.

95. Hermans D, Gautam S, García-Cañaveras JC, Gromer D, Mitra S, Spolski R, et al. Lactate dehydrogenase inhibition synergizes with IL-21 to promote CD8+ T cell stemness and antitumor immunity. Proc Natl Acad Sci U S A. Proceedings of the National Academy of Sciences; 2020;117:6047–55.

96. Liu Y-N, Yang J-F, Huang D-J, Ni H-H, Zhang C-X, Zhang L, et al. Hypoxia induces mitochondrial defect that promotes T cell exhaustion in tumor microenvironment through MYC-regulated pathways. Front Immunol. 2020;11:1906.

97. Peng J, Qin C, Ramatchandirin B, Pearah A, Guo S, Hussain M, et al. Activation of the canonical ER stress IRE1-XBP1 pathway by insulin regulates glucose and lipid metabolism. J Biol Chem. Elsevier BV; 2022;298:102283.

98. Meng H, Gonzales NM, Lonard DM, Putluri N, Zhu B, Dacso CC, et al. XBP1 links the 12-hour clock to NAFLD and regulation of membrane fluidity and lipid homeostasis. Nat Commun. Springer Science and Business Media LLC; 2020;11:6215.

99. Luo X, Alfason L, Wei M, Wu S, Kasim V. Spliced or unspliced, that is the question: The biological roles of XBP1 isoforms in pathophysiology. Int J Mol Sci. MDPI AG; 2022;23:2746.

100. Kouo T, Huang L, Pucsek AB, Cao M, Solt S, Armstrong T, et al. Galectin-3 shapes antitumor immune responses by suppressing CD8+ T cells via LAG-3 and inhibiting expansion of plasmacytoid dendritic cells. Cancer Immunol Res. American Association for Cancer Research (AACR); 2015;3:412–23.

101. Yakubovich E, Cook DP, Rodriguez GM, Vanderhyden BC. Mesenchymal ovarian cancer cells promote CD8+ T cell exhaustion through the LGALS3-LAG3 axis. NPJ Syst Biol Appl. 2023;9:61.

102. Tekguc M, Wing JB, Osaki M, Long J, Sakaguchi S. Treg-expressed CTLA-4 depletes CD80/CD86 by trogocytosis, releasing free PD-L1 on antigen-presenting cells. Proc Natl Acad Sci U S A. Proceedings of the National Academy of Sciences; 2021;118:e2023739118.

103. Hugaboom MB, Wirth LV, Street K, Ruthen N, Jegede OA, Schindler NR, et al. Presence of tertiary lymphoid structures and exhausted tissue-resident T cells determines clinical response to PD-1 blockade in renal cell carcinoma. Cancer Discov [Internet]. American Association for Cancer Research (AACR); 2025; Available from: 10.1158/2159-8290.CD-24-0991

104. Luo H, Xia X, Huang L-B, An H, Cao M, Kim GD, et al. Pan-cancer single-cell analysis reveals the heterogeneity and plasticity of cancer-associated fibroblasts in the tumor microenvironment. Nat Commun. 2022;13:6619.

105. Gao Y, Li J, Cheng W, Diao T, Liu H, Bo Y, et al. Cross-tissue human fibroblast atlas reveals myofibroblast subtypes with distinct roles in immune modulation. Cancer Cell. Elsevier BV; 2024;42:1764–1783.e10.

106. Priego N, de Pablos-Aragoneses A, Perea-García M, Pieri V, Hernández-Oliver C, Álvaro-Espinosa L, et al. TIMP1 mediates astrocyte-dependent local immunosuppression in brain metastasis acting on infiltrating CD8+ T cells. Cancer Discov. American Association for Cancer Research (AACR); 2025;15:179–201.

107. Zhang F, Wang H, Wang X, Jiang G, Liu H, Zhang G, et al. TGF-β induces M2-like macrophage polarization via SNAIL-mediated suppression of a pro-inflammatory phenotype. Oncotarget. 2016;7:52294–306.

108. Pei G, Min J, Rajapakshe KI, Branchi V, Liu Y, Selvanesan BC, et al. Spatial mapping of transcriptomic plasticity in metastatic pancreatic cancer. Nature. Springer Science and Business Media LLC; 2025;642:212–21.

109. Feig C, Jones JO, Kraman M, Wells RJB, Deonarine A, Chan DS, et al. Targeting CXCL12 from FAP-expressing carcinoma-associated fibroblasts synergizes with anti-PD-L1 immunotherapy in pancreatic cancer. Proc Natl Acad Sci U S A. Proceedings of the National Academy of Sciences; 2013;110:20212–7.

110. Portella L, Bello AM, Scala S. CXCL12 signaling in the tumor microenvironment. Adv Exp Med Biol. 2021;1302:51–70.

111. Wang Y, Szretter KJ, Vermi W, Gilfillan S, Rossini C, Cella M, et al. IL-34 is a tissue-restricted ligand of CSF1R required for the development of Langerhans cells and microglia. Nat Immunol. Springer Science and Business Media LLC; 2012;13:753–60.

112. Deshpande K, Martirosian V, Nakamura BN, Iyer M, Julian A, Eisenbarth R, et al. Neuronal exposure induces neurotransmitter signaling and synaptic mediators in tumors early in brain metastasis. Neuro Oncol. Oxford University Press (OUP); 2022;24:914–24.

113. Biermann J, Melms JC, Amin AD, Wang Y, Caprio LA, Karz A, et al. Dissecting the treatment-naive ecosystem of human melanoma brain metastasis. Cell. Elsevier BV; 2022;185:2591–2608.e30.

114. Xing X, Zhong J, Biermann J, Duan H, Zhang X, Shi Y, et al. Pan-cancer human brain metastases atlas at single-cell resolution. Cancer Cell [Internet]. Elsevier BV; 2025; Available from: 10.1016/j.ccell.2025.03.025

115. Zheng Y, Li L, Shen Z, Wang L, Niu X, Wei Y, et al. Mechanisms of neural infiltration-mediated tumor metabolic reprogramming impacting immunotherapy efficacy in non-small cell lung cancer. J Exp Clin Cancer Res. Springer Science and Business Media LLC; 2024;43:284.

116. Fernández-Nogueira P, Linzoain-Agos P, Cueto-Remacha M, De la Guia-Lopez I, Recalde-Percaz L, Parcerisas A, et al. Role of semaphorins, neuropilins and plexins in cancer progression. Cancer Lett. Elsevier BV; 2024;606:217308.

117. Evans EE, Jonason AS, Bussler H, Torno S, Veeraraghavan J, Reilly C. Antibody blockade of semaphorin 4D promotes immune infiltration into tumor and enhances response to other immunomodulatory therapies. Cancer Immunol Res American Association for Cancer Research (AACR). 2015;3:689–701.

118. Massari F, Ciccarese C, Santoni M, Lopez-Beltran A, Scarpelli M, Montironi R, et al. Targeting fibroblast growth factor receptor (FGFR) pathway in renal cell carcinoma. Expert Rev Anticancer Ther. Informa UK Limited; 2015;15:1367–9.

119. Liang P, Li Z, Chen Z, Chen Z, Jin T, He F, et al. Metabolic reprogramming of glycolysis, lipids, and amino acids in tumors: Impact on CD8+ T cell function and targeted therapeutic strategies. FASEB J. Wiley; 2025;39:e70520.

120. Xiao Y, Hu F, Chi Q. Single-cell RNA sequencing and spatial transcriptome reveal potential molecular mechanisms of lung cancer brain metastasis. Int Immunopharmacol. Elsevier BV; 2024;140:112804.

121. Ferraro GB, Ali A, Luengo A, Kodack DP, Deik A, Abbott KL, et al. Fatty acid synthesis is required for breast cancer brain metastasis. Nat Cancer. Springer Science and Business Media LLC; 2021;2:414–28.

122. Fukumura K, Malgulwar PB, Fischer GM, Hu X, Mao X, Song X, et al. Multi-omic molecular profiling reveals potentially targetable abnormalities shared across multiple histologies of brain metastasis. Acta Neuropathol. Springer Science and Business Media LLC; 2021;141:303–21.

123. Ciminera AK, Jandial R, Termini J. Metabolic advantages and vulnerabilities in brain metastases. Clin Exp Metastasis. Springer Science and Business Media LLC; 2017;34:401–10.

124. Tyagi A, Wu S-Y, Watabe K. Metabolism in the progression and metastasis of brain tumors. Cancer Lett. Elsevier BV; 2022;539:215713.

125. Venkatraman S, Balasubramanian B, Thuwajit C, Meller J, Tohtong R, Chutipongtanate S. Targeting MYC at the intersection between cancer metabolism and oncoimmunology. Front Immunol. Frontiers Media SA; 2024;15:1324045.

126. Vízkeleti L, Spisák S. Rewired metabolism caused by the oncogenic deregulation of MYC as an attractive therapeutic target in cancers. Cells [Internet]. 2023;12. Available from: 10.3390/cells12131745

127. Hoover G, Gilbert S, Curley O, Obellianne C, Lin MT, Hixson W, et al. Nerve-to-cancer transfer of mitochondria during cancer metastasis. Nature [Internet]. Springer Science and Business Media LLC; 2025; Available from: 10.1038/s41586-025-09176-8

128. Chen JC, Perez-Lorenzo R, Saenger YM, Drake CG, Christiano AM. IKZF1 enhances immune infiltrate recruitment in solid tumors and susceptibility to immunotherapy. Cell Syst. 2018;7:92–103.e4.

129. Chen E, Wu J, Huang J, Zhu W, Sun H, Wang X, et al. FLI1 promotes IFN-γ-induced kynurenine production to impair anti-tumor immunity. Nat Commun. Springer Science and Business Media LLC; 2024;15:4590.

130. Stanczak MA, Läubli H. Siglec receptors as new immune checkpoints in cancer. Mol Aspects Med. Elsevier BV; 2023;90:101112.

131. Myers KV, Amend SR, Pienta KJ. Targeting Tyro3, Axl and MerTK (TAM receptors): implications for macrophages in the tumor microenvironment. Mol Cancer [Internet]. Springer Science and Business Media LLC; 2019;18. Available from: 10.1186/s12943-019-1022-2

132. Jung K, Jeon Y-K, Jeong DH, Byun JM, Bogen B, Choi I. VSIG4-expressing tumor-associated macrophages impair anti-tumor immunity. Biochem Biophys Res Commun. Elsevier BV; 2022;628:18–24.

133. Lorenzo D, Bolli A, Tarone E, Cavallo L, Conti F. Toll-like receptor 2 at the crossroad between cancer cells, the immune system, and the Microbiota. Int J Mol Sci MDPI AG. 2020;21.

134. Sinha P, Okoro C, Foell D, Freeze HH, Ostrand-Rosenberg S, Srikrishna G. Proinflammatory S100 proteins regulate the accumulation of myeloid-derived suppressor cells. J Immunol. Oxford University Press (OUP); 2008;181:4666–75.

135. Chen Y, Ouyang Y, Li Z, Wang X, Ma J. S100A8 and S100A9 in cancer. Biochim Biophys Acta Rev Cancer. Elsevier BV; 2023;1878:188891.

136. Alban TJ, Bayik D, Otvos B, Rabljenovic A, Leng L, Jia-Shiun L, et al. Glioblastoma myeloid-derived suppressor cell subsets express differential macrophage migration inhibitory factor receptor profiles that can be targeted to reduce immune suppression. Front Immunol. Frontiers Media SA; 2020;11:1191.

137. Xu W, Dong J, Zheng Y, Zhou J, Yuan Y, Ta HM. Immune-checkpoint protein VISTA regulates antitumor immunity by controlling myeloid cell-mediated inflammation and immunosuppression. Cancer Immunol Res American Association for Cancer Research (AACR). 2019;7:1497–510.

138. Zhang S, Lv K, Liu Z, Zhao R, Li F. Fatty acid metabolism of immune cells: a new target of tumour immunotherapy. Cell Death Discov. 2024;10:39.

139. Huang J, Zhang J, Guo Z, Li C, Tan Z, Wang J, et al. Easy or not-the advances of EZH2 in regulating T cell development, differentiation, and activation in antitumor immunity. Front Immunol. Frontiers Media SA; 2021;12:741302.

140. Jiang B, Wu Z, Zhang Y, Yang X. Associations between tertiary lymphoid structure density and immune checkpoint inhibitor efficacy in solid tumors: systematic review and meta-analysis. Front Immunol. 2024;15:1414884.

141. Flavell RA, Sanjabi S, Wrzesinski SH, Licona-Limón P. The polarization of immune cells in the tumour environment by TGFbeta. Nat Rev Immunol. Springer Science and Business Media LLC; 2010;10:554–67.

142. Igarashi Y, Seino K-I. Role of IL-34 in tumors and its application to regulate inflammation. Cancer Sci. Wiley; 2025;116:1164–70.

143. Zheng GXY, Terry JM, Belgrader P, Ryvkin P, Bent ZW, Wilson R, et al. Massively parallel digital transcriptional profiling of single cells. Nat Commun. Springer Science and Business Media LLC; 2017;8:14049.

144. Schneider VA, Graves-Lindsay T, Howe K, Bouk N, Chen H-C, Kitts PA, et al. Evaluation of GRCh38 and de novo haploid genome assemblies demonstrates the enduring quality of the reference assembly. Genome Res. 2017;27:849–64.

145. Lander ES, Linton LM, Birren B, Nusbaum C, Zody MC, Baldwin J, et al. Initial sequencing and analysis of the human genome. Nature. Springer Science and Business Media LLC; 2001;409:860–921.

